# Egyptian rousette bat macrophages elicit divergent interferon responses and cytokine storm signalling against Marburg and Sudan viruses

**DOI:** 10.1101/2025.08.05.668667

**Authors:** Ivet A. Yordanova, Catherine E. Arnold, Nicolas Corrales, Jonathan C. Guito, Angelika Lander, Lay Teng Ang, Jonathan S. Towner, Joseph B. Prescott

**Affiliations:** Centre for Biological Threats and Special Pathogens, Robert Koch Institute, 13353 Berlin, Germany; Diagnostic Systems Division, US Army Medical Research Institute of Infectious Diseases, Ft. Detrick MD 21701, USA; Viral Special Pathogens Branch, Centers for Disease Control and Prevention, Atlanta, GA 30329, USA; Stanford Institute for Stem Cell Biology & Regenerative Medicine, Stanford University, Stanford, CA 94305 USA; Department of Urology, Stanford University, Stanford, USA

**Keywords:** Marburg virus, Sudan virus, filovirus, Egyptian rousette bat, reservoir, macrophage, innate immune response

## Abstract

Egyptian rousette bats (ERBs) are the only known natural reservoir of Marburg virus (MARV), etiologic agent of a highly-pathogenic zoonotic viral hemorrhagic fever. In ERBs, evolutionary adaptations allow for fine-tuned discrete pro-inflammatory immune responses that control MARV infection, yet permit population-level viral maintenance. To look for exclusive co-adapted responses between ERBs and MARV, we compared macrophage (MΦ) responses to MARV and Sudan virus (SUDV), a related filovirus not hosted by ERBs. We queried whether MARV counters normal ERB MΦ responses, illuminating co-adapted host responses not observed upon infection with SUDV, which fails to establish a productive infection and is efficiently immunologically cleared by ERBs. We observed stark differences in MΦ transcriptional responses to MARV and SUDV, including differences in Type I and III interferon (IFN)-related genes, cytokines, chemokines, cell growth and proliferation genes. We show for the first time that while MARV-infected bat MΦs undergo muted IFN responses and cytokine storm signalling, SUDV induces unperturbed Type I and III IFN gene expression, stronger cytokine and chemokine responses resembling typical host responses to a foreign viral pathogen. Our findings therefore corroborate growing evidence of unique coevolutionary relationships between bats and the specific viruses they harbor.

## Introduction

Bats possess an exceptional kaleidoscope of evolutionary adaptations, including powered flight, unique among mammals, species diversity second only to rodents, and diverse diets including but not limited to blood, nectar, fruit, insects and fish. In addition to being essential natural pollinators, bats are increasingly being recognized as important reservoirs of high-consequence viral zoonoses, including various henipaviruses, rhabdoviruses and filoviruses^1–3^. Due to their ability to transmit zoonotic pathogens to humans, domestic animals or other wildlife, several bat species represent a significant spillover risk. Hence, understanding how these zoonotic viruses have co-adapted to their natural reservoirs and are ecologically maintained is of paramount importance.

The ability of some bats to host zoonotic viruses otherwise pathogenic in humans and non-human primates (NHPs) has fostered rising interest in bat biology, ecology, viral diversity, and immune system evolution^4–10^. The findings from many of these studies illustrate that different bats have evolved diverse and highly-specific molecular mechanisms that likely contribute to their ability to tolerate and transmit viral pathogens by striking a fine-tuned balance of inducing sufficient antiviral immune responses to clear infection without aberrant tissue-damaging inflammatory processes.

Among the best characterized bat-zoonotic pathogen relationships to date is that of Egyptian rousette bats (ERBs, *Rousettus aegyptiacus*), the only verified natural reservoir of the orthomarburgviruses Marburg (MARV) and Ravn (RAVV)^11–16^. Unlike orthomarburgviruses, the natural reservoir of pathogenic orthoebolaviruses like Ebola virus (EBOV; species *Orthoebolavirus zairense*) and Sudan virus (SUDV; species *Orthoebolavirus sudanense*) remains unknown, even though bat species other than ERBs are suspected as natural reservoirs^17–20^. Along with EBOV and SUDV, MARV is an etiologic agent of sporadic outbreaks of viral hemorrhagic fever across Sub-Saharan Africa, with case fatality rates ranging from 40% to 90%^21,22^. Humans and NHPs infected with EBOV, SUDV or MARV typically develop initial non-specific flu-like symptoms, including high fever, muscle and joint pain, often followed by the rapid development of severe neurologic and hemorrhagic symptoms^23–25^. In contrast, ERBs support MARV replication in diverse tissues and shed infectious virus in the absence of signs of inflammatory disease, highlighting their reliance on a refined and highly specific co-adapted relationship with MARV^15,26,27^.

Previously, we successfully differentiated bone marrow-derived dendritic cells (bmDC) from ERBs, demonstrating that these cells support low-level MARV infection and intracellular replication. MARV-infected DCs elicited a balanced response involving upregulated canonical antiviral signalling genes and suppressed proinflammatory cytokine/chemokine gene expression^28^. Similarly, CD14^+^ monocyte-like cells isolated from the spleens of MARV-infected ERBs support viral replication and display transient upregulation of genes associated with Type I IFN responses, viral restriction, and anti-inflammatory signalling pathways^29^. In the liver, MARV-infected ERBs harbor discrete foci of inflammation in the absence of notable tissue pathology elsewhere^27^. In infected bats, viral RNA is clearly detectable in Iba1^+^ mononuclear phagocytes in the liver, as well as in follicular DC-like cells in axillary lymph nodes, underlining the specific cell tropism of MARV for host MΦs and DCs^27^. Similar host cell tropism is also observed in humans and NHPs, where filoviruses induce significant tissue-damaging proinflammatory cytokine and chemokine release in MΦs, while DCs undergo arrested maturation and display dysregulated antigen presentation functions^30–36^.

Contrasting their reservoir competence for MARV, ERBs are generally refractory to orthoebolaviruses like EBOV, Bundibugyo virus (BDBV), Taï Forest virus (TAFV) or Reston virus (RESTV)^26^. Only low levels of SUDV viral RNA have been detected in select tissues of experimentally infected ERBs, in the absence of viral shedding. Comparative analysis of MARV and SUDV viral loads in liver, spleen and kidney tissues shows ERBs control SUDV replication faster than MARV, indicating differential control of the two filoviruses^26^. However, the underlying innate immune mechanisms potentially contributing to this divergent control of filovirus infections in these bats remain unknown. Directly comparing ERB MΦ responses to MARV and SUDV infections therefore offers an invaluable opportunity to elucidate specific host responses that have evolved during co-adaptation that allow for the development of a productive MARV infection, but a non-productive “dead-end” infection with SUDV. By leveraging our ability to generate bone marrow-derived MΦs, we were able to show that MARV evades specific features of ERB immunity, including macrophage activation and type III IFN responses, both of which are induced by SUDV, likely contributing to the ability of these bats to efficiently combat SUDV. In contrast, MARV is able to evade immunity to prolong replication and infection by relying on a complex combination of viral protein antagonism patterns and host cell cytoskeletal changes unique to ERBs.

## Results

### ERB bone marrow cells differentiate into MΦs in response to recombinant bat M-CSF

Dysregulation of host MΦs is a classical feature of filovirus disease. We therefore sought to examine the immune response profile of filovirus-infected ERB MΦs. For this, we optimized our existing approach for ERB bmDC differentiation into a novel protocol to reproducibly generate bmMΦs *in vitro* (**Fig. 1a**). Bone marrow cells isolated from naïve bats were cultured in medium containing ERB-specific recombinant macrophage colony-stimulating factor (rM-CSF; Kingfisher Biotech). In the presence of rM-CSF, bone marrow cells consistently developed heterogeneous morphology after 8 days, but maintained consistent adherence properties and displayed dendrites, typical morphological features of macrophages (**Fig. 1b, Fig. S1a**). In contrast, cultures without rM-CSF contained only small, non-adherent cell-like particles and debris (**Fig. 1b**).

**Figure 1.**
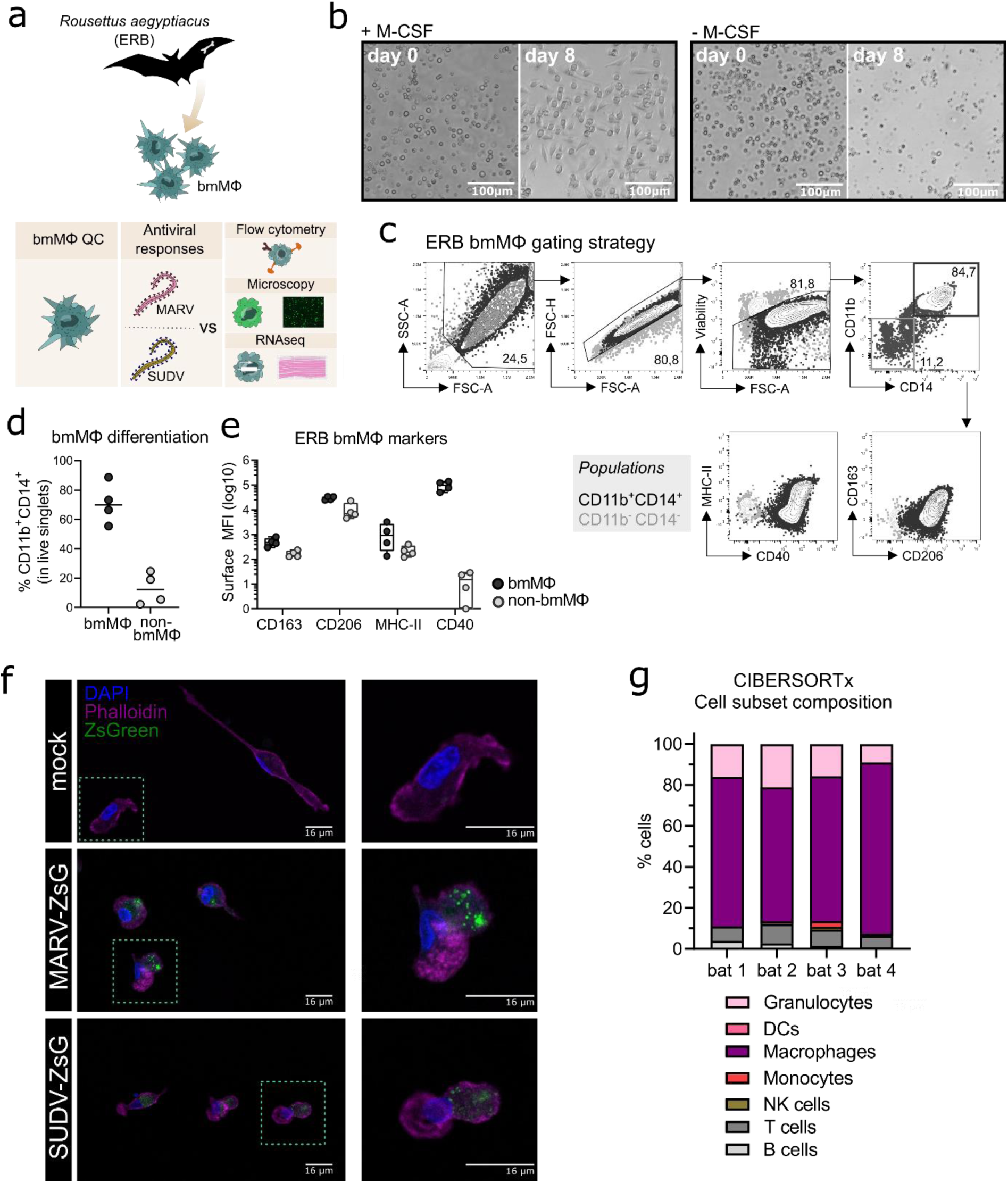
Differentiation and initial characterization of Egyptian rousette bat (ERB) bone marrow-derived macrophages (bmMΦs). (a) Graphical summary of the experimental framework. (b) Example brightfield microscopy images illustrating the morphology of bone marrow cell cultures following 8 days of *in vitro* differentiation with (left) or without (right) recombinant ERB macrophage colony-stimulating factor (M-CSF). (c) Gating strategy and example contour plots showing the identification of CD11b^+^CD14^+^ bmMΦs (black gate) via flow cytometry and their expression of surface markers CD40, MHC-II, CD163 and CD206, overlaid with the expression of the same markers in undifferentiated CD11b^-^CD14^-^ cells (grey gate). (d) Percentages of CD11b^+^CD14^+^ and CD11b^-^CD14^-^ cells among live singlets in M-CSF-differentiated bone marrow cultures. (e) Median fluorescence intensities (MFI) of surface markers CD163, CD206, MHC-II and CD40 expressed on CD11b^+^CD14^+^ bmMΦ and CD11b^-^ CD14^-^ non-bmMΦ cells after 8 days of culture. (f) Representative confocal microscopy images of mock, MARV-ZsG and SUDV-ZsG-infected bmMΦs. Scale bars correspond to 16µm. Cell nuclei are stained in blue (DAPI) and actin filaments in magenta (Phalloidin). Virus-infected cells contain fluorescent ZsGreen signal. The smaller panels on the right illustrate zoomed in images of the cells marked with dotted squares in the main panels on the left. (g) CIBERSORTx analysis of the baseline cell subset composition of ERB-derived bmMΦs. The analysis used the transcriptional profile of mock-infected bmMΦs compared against known signatures identified for human marker genes using a software-defined signature matrix to calculate the proportions of cell types. The results in (d) and (e) are pooled from two independent experiments with n=2 bats per experiment.

To confirm the phenotype of our ERB bmMΦs, we quantified the cell surface expression of canonical MΦ markers, including myeloid cell markers CD11b and CD14, the scavenger receptor CD163, the mannose receptor CD206, the antigen presentation receptor MHC-II and the costimulatory marker CD40 (**Fig. 1c-e**). Bone marrow cells cultured with rM-CSF generated variable proportions of bmMΦs from individual bats, averaging 65-85% of myeloid CD11b^+^CD14^+^ cells (**Fig. 1d; Fig. S1b-e**). Contrasting with CD11b^-^CD14^-^ cells (non-bmMΦ), the CD11b^+^CD14^+^ population (bmMΦ) had higher surface expression of CD163, CD206 and MHC-II, while CD40 expression in bmMΦs was almost 10^4^-fold higher than in non-bmMΦ (**Fig. 1e**).

Host MΦs are among the first targets of filoviruses *in vivo*. To assess whether ERB-derived bmMΦs are susceptible to MARV and SUDV, and to better capture their morphology at baseline and following initial filovirus infection, bmMΦs were infected with recombinant MARV or SUDV viruses expressing a green fluorescent protein (ZsGreen, ZsG) or were left uninfected. Using high-resolution confocal microscopy and staining of cell nuclei (DAPI, blue) and the cytoskeleton (Phalloidin, magenta), we observed classic macrophage morphology (**Fig. 1f**). In MARV-infected cells we observed granular foci of strong ZsG signal, possibly as a result of partially incomplete cleavage of the NP-ZsG fusion protein as previously reported^37^. The cells were also readily susceptible to SUDV, evidenced by the presence of strong but more diffuse cytoplasmic ZsG signal in line with its VP40-ZsG fusion protein construct and typical assembly mechanism of SUDV in host cells^38,39^. Together, these microscopy findings illustrate that ERB-derived bmMΦs display a classical macrophage morphology and demonstrate evidence of characteristic viral replication and assembly.

Finally, we performed bulk RNA sequencing of freshly-differentiated bmMΦs to assess their baseline transcriptional profile as an additional quality control step. Using the complete transcriptional profile of the cells, we assessed their cell culture composition using CIBERSORTx and could show that our cultures were predominantly classed as “macrophages” based on their complete gene expression profile (**Fig. 1g**).

### MARV replication dynamics differ from SUDV in ERB innate immune cells

To detect differences in viral transcription efficiency between MARV and SUDV, ERB bmMΦs were infected with wild-type MARV or SUDV, or recombinant fluorescent ZsG viruses (MARV-ZsG or SUDV-ZsG) to monitor the kinetics of viral infection, replication and progeny production. Cells were infected with each virus at a multiplicity of infection (MOI) of 2 (measured on Vero E6 cells), and samples were collected over 3 days for RNA sequencing and qRT-PCR (wild-type viruses) or were observed microscopically (ZsG-expressing viruses).

Cells infected with either virus displayed clear signs of viral protein transcription over the course of the 3-day infection, evidenced by the presence of ZsG signal in virus-infected bmMΦ cultures (**Fig. 2a**). Bulk RNAseq analysis revealed that SUDV-infected cells harbored higher intracellular viral gene copy numbers of NP (3.7-fold higher), VP35 (4.8-fold higher), VP40 (2.5-fold higher) and GP (7-fold higher) compared with MARV-infected cells, indicative of higher viral replication of SUDV in these cells and in line with the stronger ZsG signal observed (**Fig. 2a, b, d**). Surveying viral progeny production in cell culture supernatants, cells from individual bats maintained overall stable numbers of MARV-NP gene copies/µL supernatant between 1-3 DPI. In contrast, SUDV-infected cells showed a trend for decreasing viral progeny production between 1 DPI and 3 DPI, suggestive of efficient control of infection compared with MARV (**Fig. 2c, e**).

**Figure 2.**
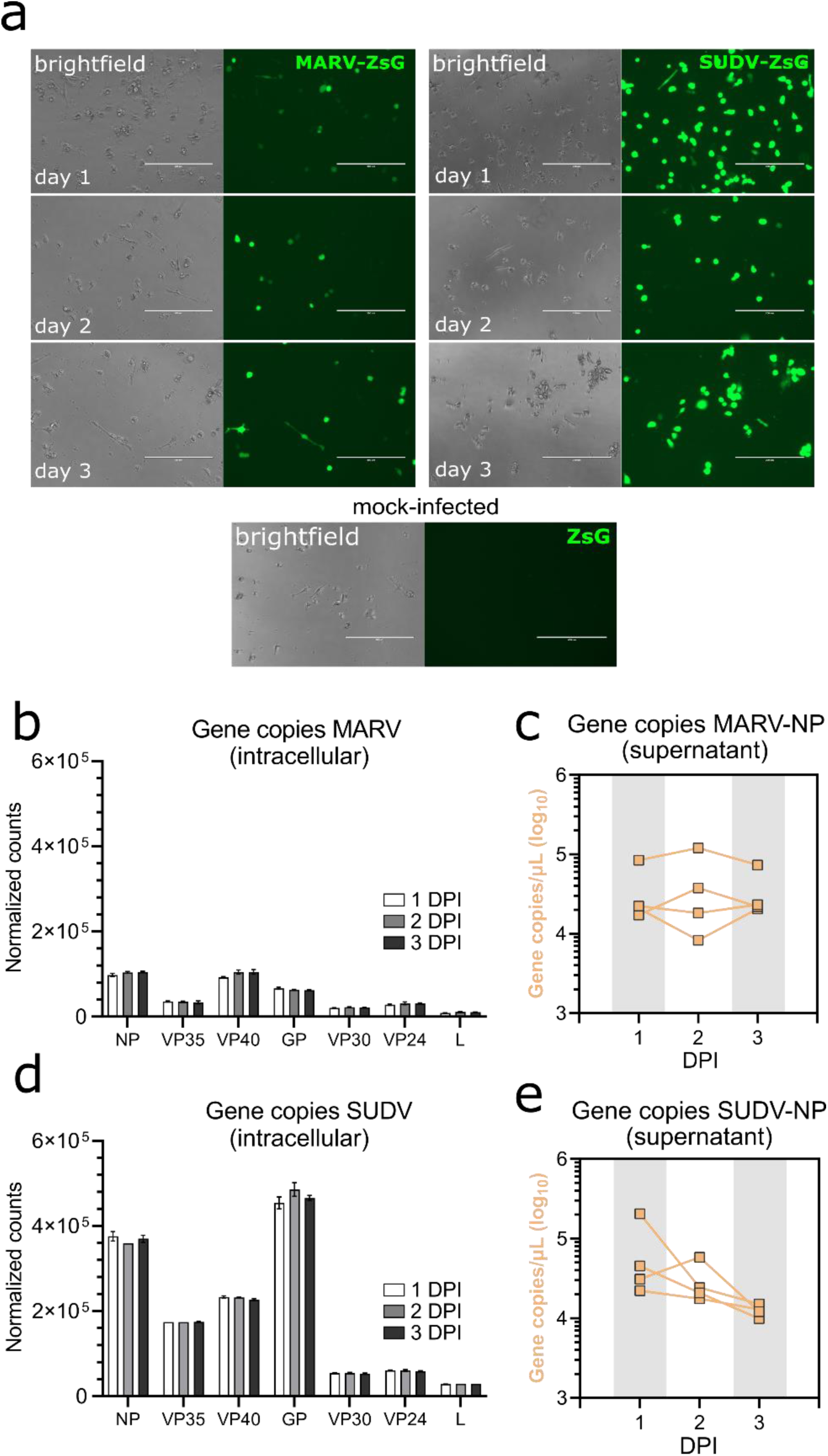
Intracellular virus replication and progeny production in ERB bmMΦs. (a) Fluorescent ZsGreen expression in MARV-ZsG-infected, SUDV-ZsG-infected and mock-infected ERB bmMΦs at 1, 2, and 3 DPI. Scale bars correspond to 200µm. (b) Normalized counts of MARV genes at 1, 2 and 3 DPI in bmMΦs. (c) Gene copy counts of MARV-NP in cell culture supernatants. (d) Normalized counts of SUDV genes at 1, 2 and 3 DPI in bmMΦs. (e) Gene copy counts of SUDV-NP in cell culture supernatants. The data in (b-e) are pooled from two independent experiments with n=2 bats per experiment.

Considering the difference in virus replication between MARV and SUDV in ERB bmMΦs, next we sought to test whether intrinsic differences in replication between the two viruses explain this observation. For this, we infected ERB kidney-derived immortalized RoNi cells and Vero E6 cells with MARV-ZsG and SUDV-ZsG. At 1, 2 and 3 DPI each cell line was surveyed via flow cytometry and qRT-PCR to quantify the percentage ZsG-positive cells and viral RNA in cell culture supernatants (**Fig. S2a-d**). Overall, MARV-ZsG and SUDV-ZsG replicated similarly in Vero E6 cells. In contrast, we observed significantly fewer ZsG^+^ cells at 2-3 DPI in MARV-infected RoNi cells, diverging from the SUDV-ZsG replication in RoNi cultures, which was comparable to that observed in VeroE6 cells (**Fig. S2a-c**). Together, these findings highlight that instead of virus-intrinsic differences, the differential replication of MARV and SUDV is host-intrinsic, likely as a result of specific co-evolutionary adaptations between ERBs and MARV.

### ERB bmMΦs mount transcriptionally distinct responses to general immune stimulation and filoviruses

Next, we applied bulk RNA sequencing to profile the transcriptional responses of bmMΦs to both general stimulation and filovirus infections, expanding the depth and breadth of our understanding of ERB innate immune responses at the cellular level. Similar to bmDCs^28^, ERB-derived bmMΦs displayed a clear transcriptional response distinct between general immune agonists like bacterial lipopolysaccharide (LPS) and Sendai virus (SeV), and the two filoviruses (**Fig. 3a**). MARV induced the differential expression of overall smaller clusters of genes than SUDV, mostly at 1 DPI and 2 DPI, while SUDV induced the consistent differential expression of larger gene sets throughout all three timepoints (**Fig. 3a**). In response to LPS, bmMΦs upregulated various transcriptional factors (*STAT4*), proinflammatory cytokines (*TNF, IL1A, IL6, IL12B, IL23)*, cell migration receptors (*CCR7, ITGB8*) and chemokines (*CXCL6, CCL22*) (**Fig. S3a**). In contrast, SeV infection induced canonical antiviral IFN-associated genes like *IFNB1, ISG20* and *IFIT3*, several chemokine and chemokine receptor genes (*CCL5, CXCL11, CCR3, CCR7*), proinflammatory cytokines (*IL6*) and activation markers (*CD82, CD163, CD207*) (**Fig. S3b**).

**Figure 3.**
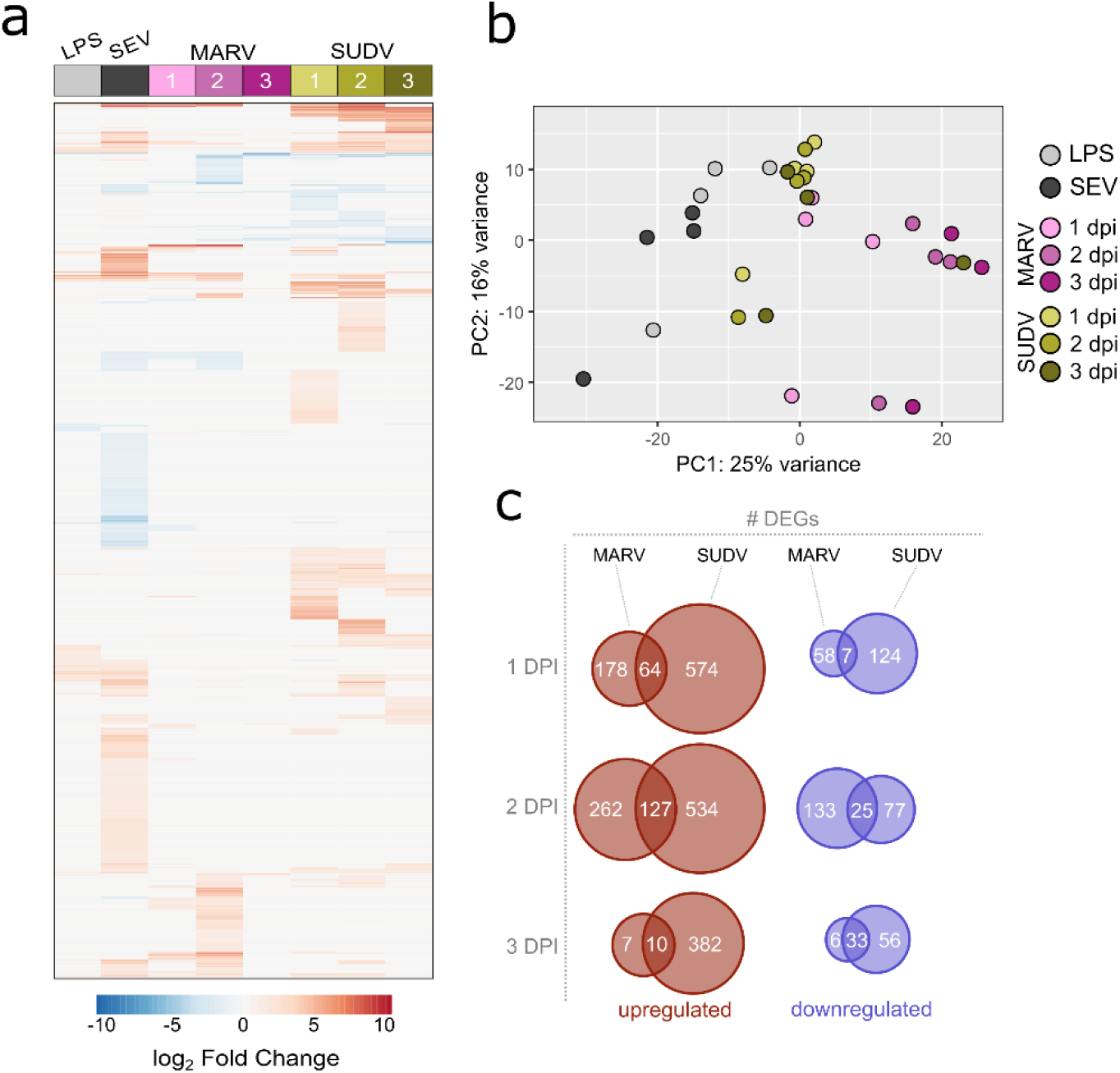
Global transcriptional responses of bat bmMΦs to stimulation and viral infection. (a) Global heatmap of differentially expressed genes (DEG) in ERB bmMΦs after 1 day of LPS restimulation and SeV infection, and at 1, 2 and 3 DPI with MARV or SUDV. DEGs were defined as genes with a padj < 0.05 and a log_2_ fold change ≥ ± 1.5. (b) Principal component analysis (PCA) plot of LPS-treated, SeV-infected and filovirus-infected bmMΦs, based on the expression levels of the top 500 most highly-expressed genes in each treatment group. (c) Venn diagram of the total numbers of significantly upregulated (red) and downregulated (blue) DEGs identified uniquely for MARV, SUDV or shared in both viral infections at 1, 2 and 3 DPI. The heatmap in (a) is shown as log_2_ fold change against mock-infected negative controls.

Principal component analysis based on the top 500 most highly differentially expressed genes confirmed the treatment and infection-specific responses across the four individual bats (**Fig. 3b**). ERB bmMΦs clustered predominantly by treatment (LPS) or virus infection (SeV/MARV/SUDV), with temporal differences in clustering mostly evident for MARV-infected samples, while most SUDV samples clustered together independent of day of infection, in line with the continuous differential expression of large gene sets shown in the DEG heatmap (**Fig. 3a, b**). SUDV induced almost 5-fold more unique upregulated DEGs (574 vs 178) and twice the number of unique downregulated differentially expressed genes (DEGs) (124 vs 58) than MARV as early as 1 DPI. While the expansion of both unique and shared DEGs for MARV was only transient and significantly contracted by 3 DPI, SUDV-infected bmMΦs maintained stable differential expression of large sets of both upregulated and downregulated DEGs unique to SUDV throughout the 3-day infection (**Fig. 3c**).

### bmMΦs initiate disparate host cell transcriptional responses to MARV and SUDV

Significant IFN-associated host cell transcriptional responses to both filoviruses consisted almost exclusively of upregulated genes, whose differential expression was mostly limited to 1-2 DPI for MARV, and 1-2 DPI or 2-3 DPI for SUDV. Within the first two days of infection with either filovirus, ERB bmMΦs upregulated an identical cluster of Type I IFN genes, including *IFNB1*, two *IFNA*-like, an *IFNA4*-like and an *IFNW1*-like gene – a response mirrored in SeV-infected cells. Another *IFNW1*-like gene and two type III IFN genes (*IFNL1*-like and *IFNL3*) were upregulated only in response to SeV and SUDV at 2 and 3 DPI (**Fig. 4a**). Beyond Type I IFNs, MARV infection induced overall muted IFN-associated gene expression in bmMΦs, characterized mostly by transient upregulation of *ISG20* and *IRF4* at 1 and 2 DPI, and the upregulated expression of four TRIM protein-coding genes at 2 DPI (*TRIM16, TRIM54, TRIM66* and *TRIM72*). SUDV induced a stronger shift in gene expression, including the stable upregulation of *STAT4* and *IFITM10* between 1-3 DPI, a delayed upregulation of multiple ISGs (*IFIH1, IFIT2, IFIT3, OASL, OAS3, ISG15, ISG20, ZBP1)*, but only transient upregulation of *TRIM66* and *TRIM72* at 2 DPI (**Fig. 4a**).

**Figure 4.**
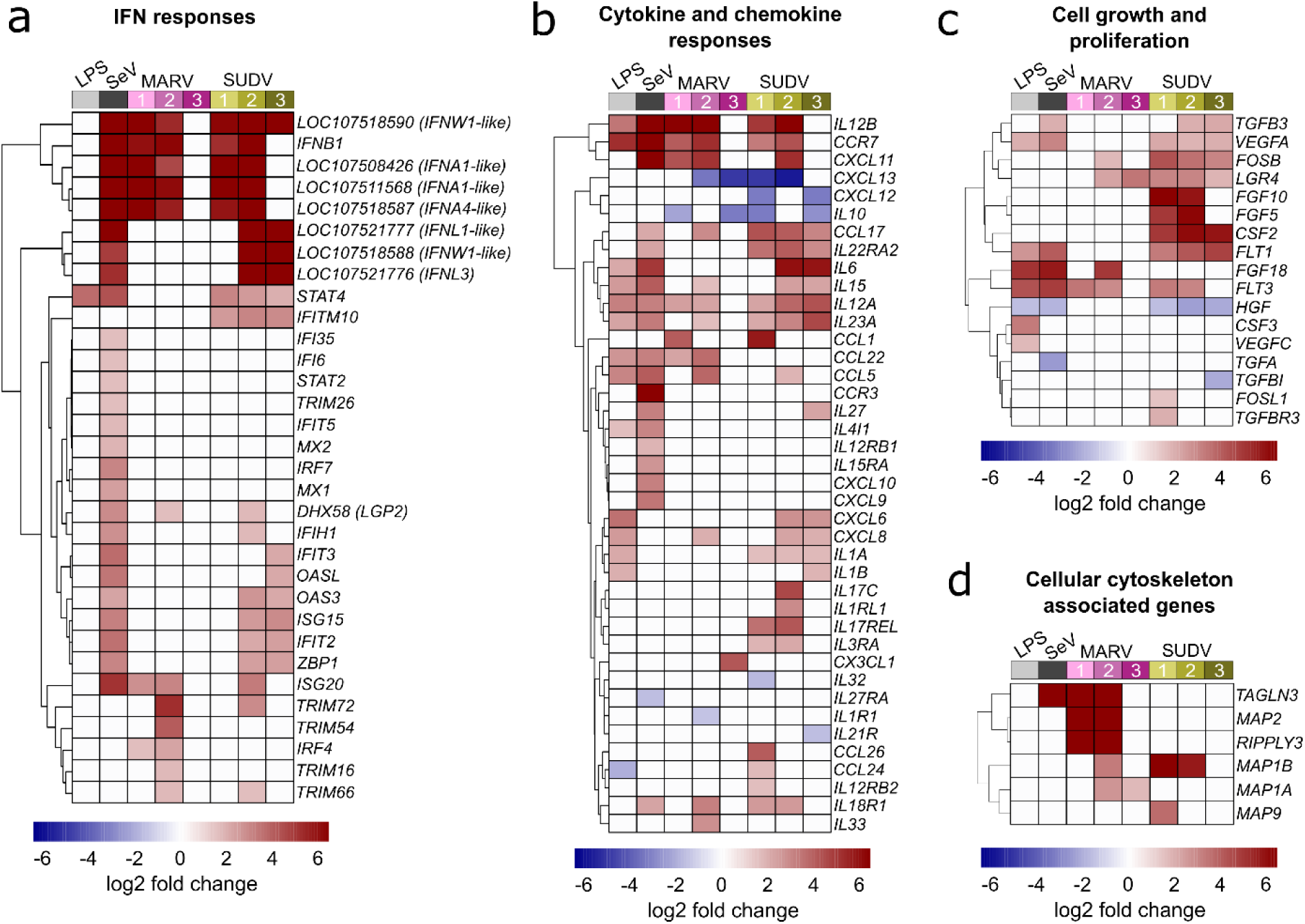
Immune-related gene expression profile of ERB-derived bmMΦs. Heatmaps of (a) IFN response DEGs; (b) Cytokine and chemokine DEGs; (c) Cell growth and proliferation-related DEGs and (d) Host cell cytoskeleton-associated DEGs in ERB bmMΦs after 1 day of LPS restimulation and SeV infection, and at 1, 2 and 3 DPI with MARV or SUDV. DEGs were defined as genes with a padj < 0.05 and a log_2_ fold change ≥ ± 1.5. Each treatment group or timepoint includes pooled data from four individual bats. Each heatmap is shown as log_2_ fold change against mock-infected negative controls.

Alongside antiviral IFN responses, ERB bmMΦs displayed diverse differential gene expression of various cytokines and chemokines in response to both filoviruses. Between 1 and 2 DPI, both MARV and SUDV upregulated the expression of *IL12B, CCR7* and *CXCL11*, reflecting a similar response in LPS-stimulated and SeV-infected cells. In contrast, the anti-inflammatory cytokine *IL10*, and the chemokines *CXCL12* and *CXCL13* were strongly downregulated in response to both MARV and SUDV, but remained unchanged following LPS stimulation or SeV infection, indicative of filovirus-specific gene suppression (**Fig. 4b**). MARV infection induced a muted sporadic upregulation at either 1 DPI or 2 DPI of several chemokines (*CCL1, CCL5, CCL17, CCL22, CXCL8, CXCL11*) and cytokines (*IL12A, IL15, IL23A, IL33*). In contrast, SUDV infection upregulated more sustained expression of *CCL17*, paralleled by a transient upregulation of more proinflammatory cytokines and chemokines at either 1-2 DPI (*CCL1, IL17C, CCL24, CCL26*) or 2-3 DPI (*IL6, IL15, IL12A, IL23A, CXCL6, CXCL8*) (**Fig. 4b**).

In parallel with the observed expression profiles of immune-related genes, filovirus-infected bmMΦs shifted their expression of several cell growth and proliferation-associated genes. SUDV induced the sustained upregulation of genes like *FOSB, LGR4, FGF5, FGF10, CSF2, FLT1* and *FLT3*. In contrast, MARV-infected cells underwent weaker transient upregulation of *FOSB*, *LGR4*, *FGF18* and *FLT3*. Interestingly, the hepatocyte growth factor *HGF*, whose gene expression was significantly downregulated in LPS-treated, SeV-infected and SUDV-infected cells remained unchanged in response to MARV, indicating potential MARV-specific regulation of growth factor expression in bat innate immune cells, potentially to limit hepatic injury. (**Fig. 4c**).

Viruses rely on the host cell machinery not only for viral genome replication, but also for trafficking to sites of replication, shuttling intermediate viral products between sites of assembly within the cytoplasm and egress of newly assembled viral particles. Considering the pronounced differences in MARV and SUDV replication in ERB bmMΦs, we next surveyed the top DEGs in filovirus-infected cells for genes associated with host cell actin filament organization and polymerase activity. Among these, we found four genes encoding microtubule associated proteins (*MAP*s), *RIPPLY3* encoding RNA polymerase II, and *TAGLN3* associated with actin filament organization and RNA polymerase II transcription. MARV infection upregulated the expression of five of these DEGs (*MAP1A, MAP1B, MAP2, RIPPLY3* and *TAGLN3*), while SUDV only induced the upregulation of *MAP1B* and a transient increase of *MAP9* at 1 DPI. Considering the sustained low-level replication of MARV, these findings could indicate virus-induced alterations of the ERB cell cytoskeleton that contribute to a slower but more sustained viral replication and progeny production that ultimately translates in the ability of wild ERBs to maintain low-level MARV infections long enough for virus transmission to other bats and long-term maintenance at the population level year-long (**Fig. 4d**). In contrast, SUDV could be maladapted to utilize the ERB intracellular machinery for efficient virus progeny assembly, trafficking and egress.

### ERB bmMΦs experience differential antagonism by filovirus-encoded proteins

Filoviruses have evolved various strategies to either evade or antagonize host innate immunity through distinct mechanisms driven mostly by filoviral VP24, VP35 and VP40 proteins. Considering the pronounced differences in transcriptional responses to MARV and SUDV in ERB-derived bmMΦs shown here, we explored whether the two viruses potentially exert differing IFN antagonistic properties, which could be associated with the distinct intracellular replication and viral progeny dynamics in these cells. For this, we used Spearman’s correlation analysis of the normalized gene counts of DEGs encoding key pattern-recognition receptor genes (*RIG-I* and *LGP2*), Type I and Type III IFNs (*IFNAs, IFNWs* and *IFNL*s), IFN receptors (*IFNAR1/2, IFNLR1* and *IL10RB*), transcriptional factors (*IRF1/3/4/7* and *NFKB1*) and interferon-stimulated genes (ISGs) (*ISG15/20*) against the two major antagonistic proteins of MARV (MARV-VP35 and MARV-VP40) and SUDV (SUDV-VP35 and SUDV-VP24), known to interfere with host IFN production, IFN-induced STAT signaling and phosphorylation, as well as RIG-I signaling.

Correlation analysis revealed that MARV-VP35 gene expression showed a moderate negative correlation with only two of the 21 surveyed ERB IFN-associated genes – *IFNB1* (ρ=-0.52) and the Type III IFN receptor *IFNLR1* (ρ=-0.61), indicative of severely limited VP35-driven suppression (**Fig. 5a**). In stark contrast, SUDV-VP35 demonstrated a moderate to strong negative correlation with all but three surveyed IFN signaling-associated genes included in the analysis, suggestive of strong SUDV-VP35-driven antagonism of ERB IFN responses (**Fig. 5b**). Unlike the negative correlation observed between MARV-VP35 and *IFNLR1*, SUDV-VP35 showed a moderate positive correlation with *IFNLR1*, in line with our earlier observation of differential induction of Type III IFNs by SUDV but not MARV. Beyond the lack of negative correlation between MARV-VP35 and ERB IFN-associated genes, however, we found that MARV-VP40 exhibited stronger negative correlation with *DDX58* (RIG-I), two IFNA-like genes, *IFNL3, IFNAR2, IL10RB, IRF7, NFKB1, ISG15* and *ISG20* than either SUDV-VP35 or SUDV-VP24, indicative of strong MARV-VP40 antagonism and hence of major differences in how each virus potentially blocks or counteracts macrophage IFN responses in ERBs (**Fig. 5a, b**).

**Figure 5.**
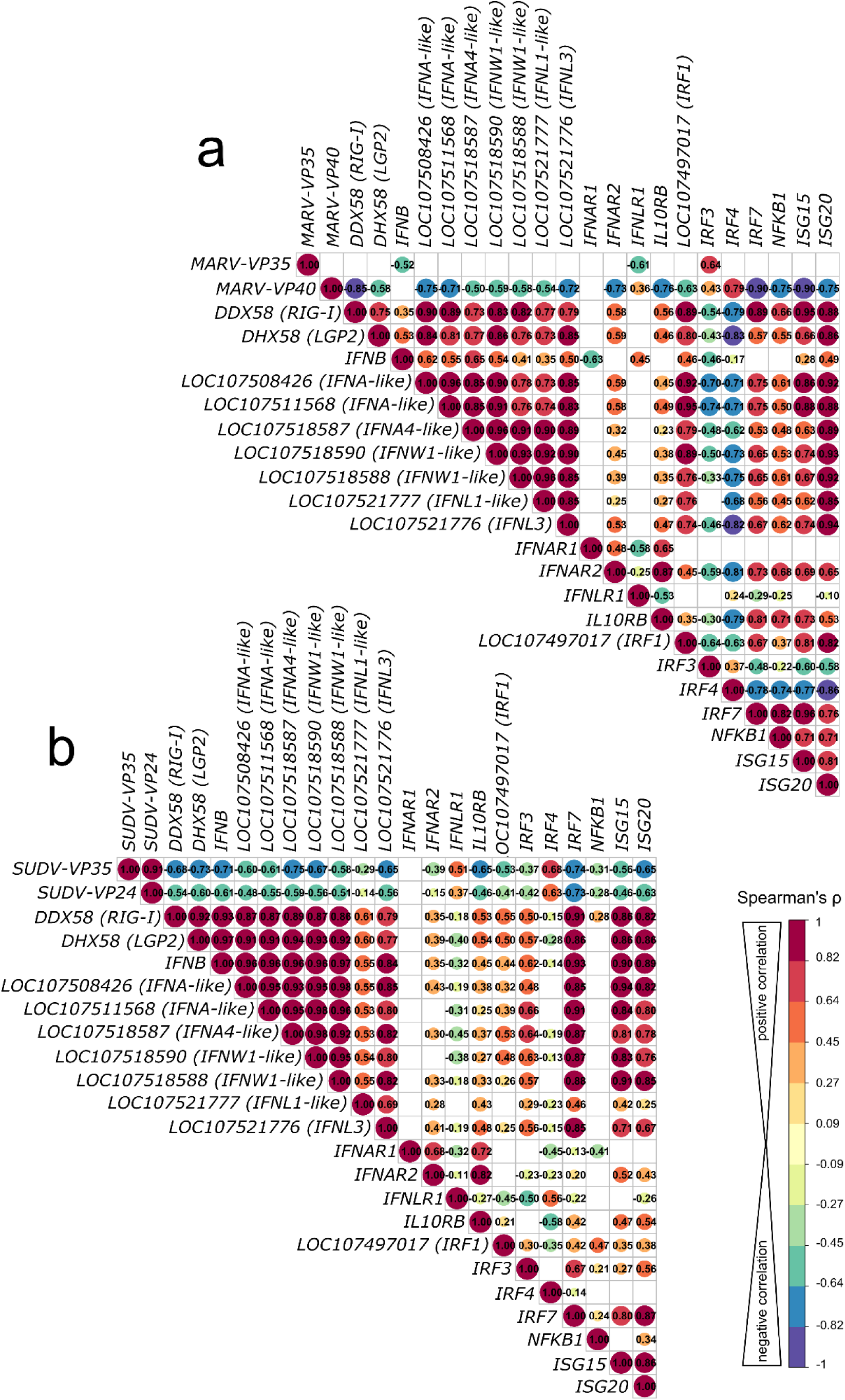
Correlation analysis of antagonistic filovirus gene and host antiviral gene expression patterns in bat bmMΦs. (a) Correlation matrix comparing correlations of gene expression of MARV infection-induced IFN response DEGs with MARV-VP35 and MARV-VP40. (b) Correlation matrix comparing correlations of gene expression of SUDV infection-induced IFN response DEGs with SUDV-VP35 and SUDV-VP24. The correlation matrix analysis used normalized gene counts of each gene as input. The data were analyzed using Spearman’s ρ correlation test using the corrr RStudio package. A perfect positive correlation is considered as having a ρ=1 and a perfect negative correlation a ρ=-1. Statistically insignificant correlations (p>0.05) are masked in the correlation matrices as empty boxes. The data used for the correlation analysis are pooled from four individual bats at 1, 2 and 3 DPI with each respective filovirus.

### MARV and SUDV elicit distinct signaling pathways in bat bmMΦs

Using Ingenuity Pathway Analysis (IPA), next we predicted what canonical signaling pathways are differentially regulated by the two filoviruses in bat-derived bmMΦs. Supplying the observed differential gene expression values for each virus at each day post-infection, the IPA software simulates the directional consequences of downstream molecules, infers upstream activity within given signaling pathways and predicts what upstream regulators may be causing observed gene expression changes and whether any canonical signaling pathways or biological processes are differentially regulated. Among the top 5 canonical signaling pathways regulated uniquely by MARV, we found two upregulated (*Mitotic G1 phase* and *G1/S transition and Senescence*) and three downregulated (*Cell Cycle Checkpoints, Synthesis of DNA* and *Cell Cycle Control of Chromosomal Replication*) pathways (**Fig. 6a**). In contrast, the top 5 canonical pathways regulated uniquely by SUDV comprised only upregulated pathways, including *Systemic Lupus Erythematosus in B cell Signaling Pathway, Hepatic Cholestasis, Dendritic Cell Maturation, Synaptogenesis Signaling Pathway* and *Pancreatic Secretion Signaling Pathway* (**Fig. 6b**). Among the top 5 shared canonical pathways regulated by both viruses were *Cardiac Hypertrophy Signaling (Enhanced), S100 Family Signaling Pathway, Macrophage Classical Activation Signaling Pathway, Pathogen Induced Cytokine Storm Signaling Pathway* and *IL-17 Signaling*, all of which were either downregulated or not at all regulated in MARV-infected cells by 3 DPI, but remained strongly upregulated in response to SUDV at all three timepoints (**Fig. 6c**).

**Figure 6.**
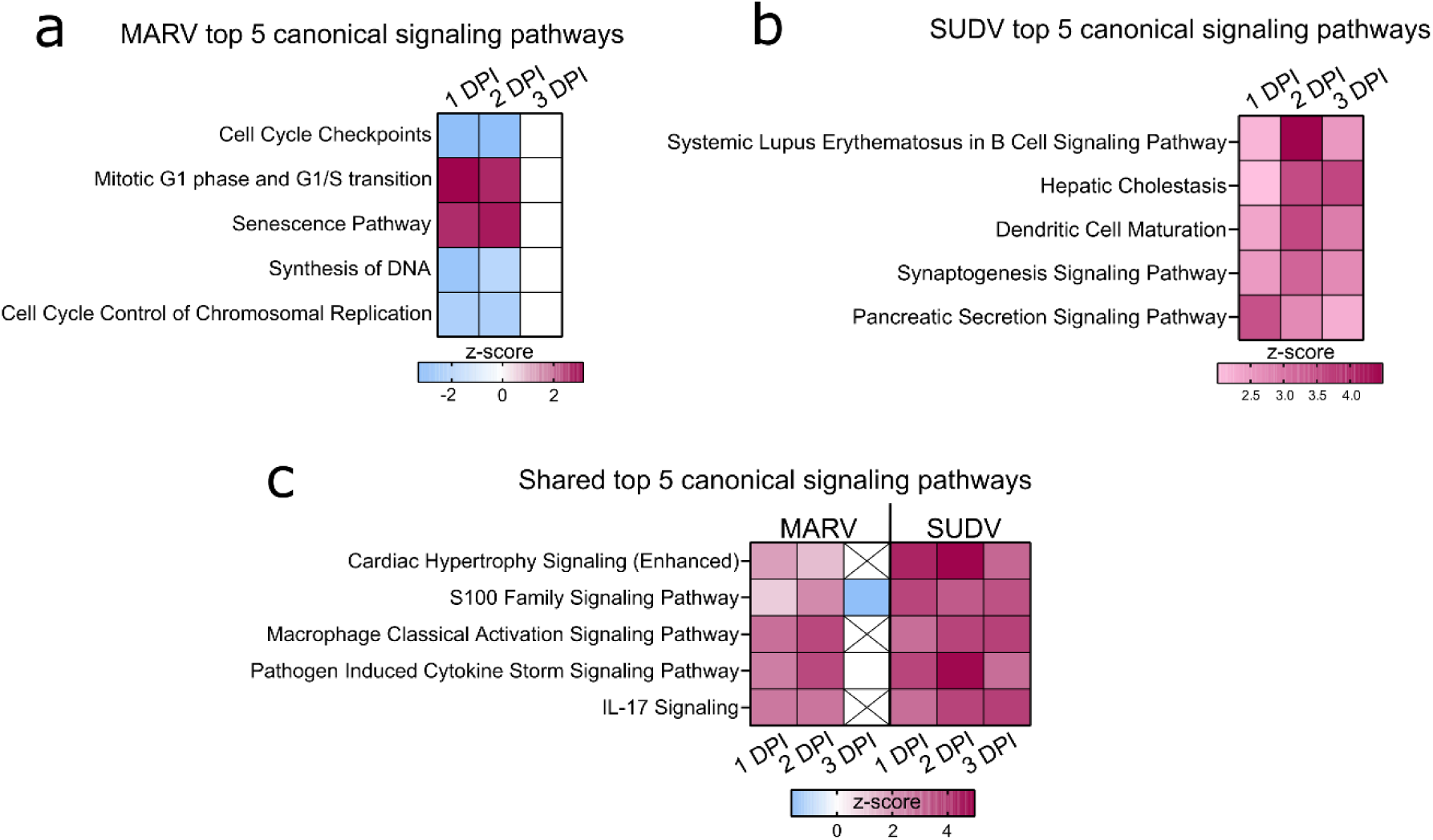
Differential regulation of canonical signaling pathways in filovirus-infected ERB bmMΦs. Ingenuity Pathway Analysis (IPA) of the top 5 statistically significant (p<0.05) canonical signaling pathways in response to (a) MARV, (b) SUDV and (c) shared by both viruses at 1, 2 and 3 DPI, illustrated as z-score heatmaps.

Next, we predicted what upstream regulators, molecules capable of regulating the expression, transcription or phosphorylation of other molecules, were differentially regulated in response to either MARV or SUDV infection. The top 20 IPA-predicted upstream regulators positively activated in response to MARV included various innate immune and proinflammatory response genes such as *TNF, IL1B, IFNG, CD40LG, TLR3* as well as *RELA,* an NFκB signaling-associated transcriptional factor. This response was largely limited to 1 DPI and 2 DPI for MARV, while at 3 DPI the top positively regulated upstream regulators only included *TNF* and several growth factors (*HGF, VEGF* and *EGF*) (**Fig. S4a**). In contrast, *IL10* encoding the canonical anti-inflammatory cytokine was negatively regulated consistently at all three timepoints in MARV-infected cells. At 3 DPI, MARV infection additionally resulted in the parallel inhibition of both *IL21* and *IL6*, two genes encoding a key immune regulatory cytokine and a canonical proinflammatory cytokine, respectively, indicating the orchestration of a carefully balanced immune response to MARV (**Fig. S4a**).

For SUDV, the top 20 positively regulated IPA-predicted upstream regulators included a mix of innate immune genes, transcriptional and growth factors. Reflecting some of the findings for MARV, SUDV-induced upstream regulators included *TNF, IL1B, CD40LG, RELA, IFNG* and *TLR3*. However, additional upstream regulators were also positively regulated in response to SUDV, such as *IRF1, IRF7*, *poly rI:rC RNA* and *NFKB (complex)*, as well as growth factors *VEGF, NGF* and *EGF* (**Fig. S4b**).

### Pathogen-induced responses in ERB bmMΦs suggest divergent regulation of downstream immune signaling

Considering the clear differences in transcriptional responses to MARV and SUDV in ERB bmMΦs, next we focused on surveying in more detail any discrepancies in the regulation of genes comprising the *Pathogen Induced Cytokine Storm Signaling Pathway*, predicted by IPA as differentially regulated by both viruses. Using IPA’s Molecular Activity Predictor (MAP) tool, we explored signaling cascades observed and predicted by MAP as differentially regulated in macrophages, endothelial cells, hepatocytes and various T cell subsets. We directly compared the gene expression regulation by MARV and SUDV, choosing the 2 DPI timepoint to survey the peak transcriptional response changes in responses to both viruses (**Fig.7**).

**Figure 7.**
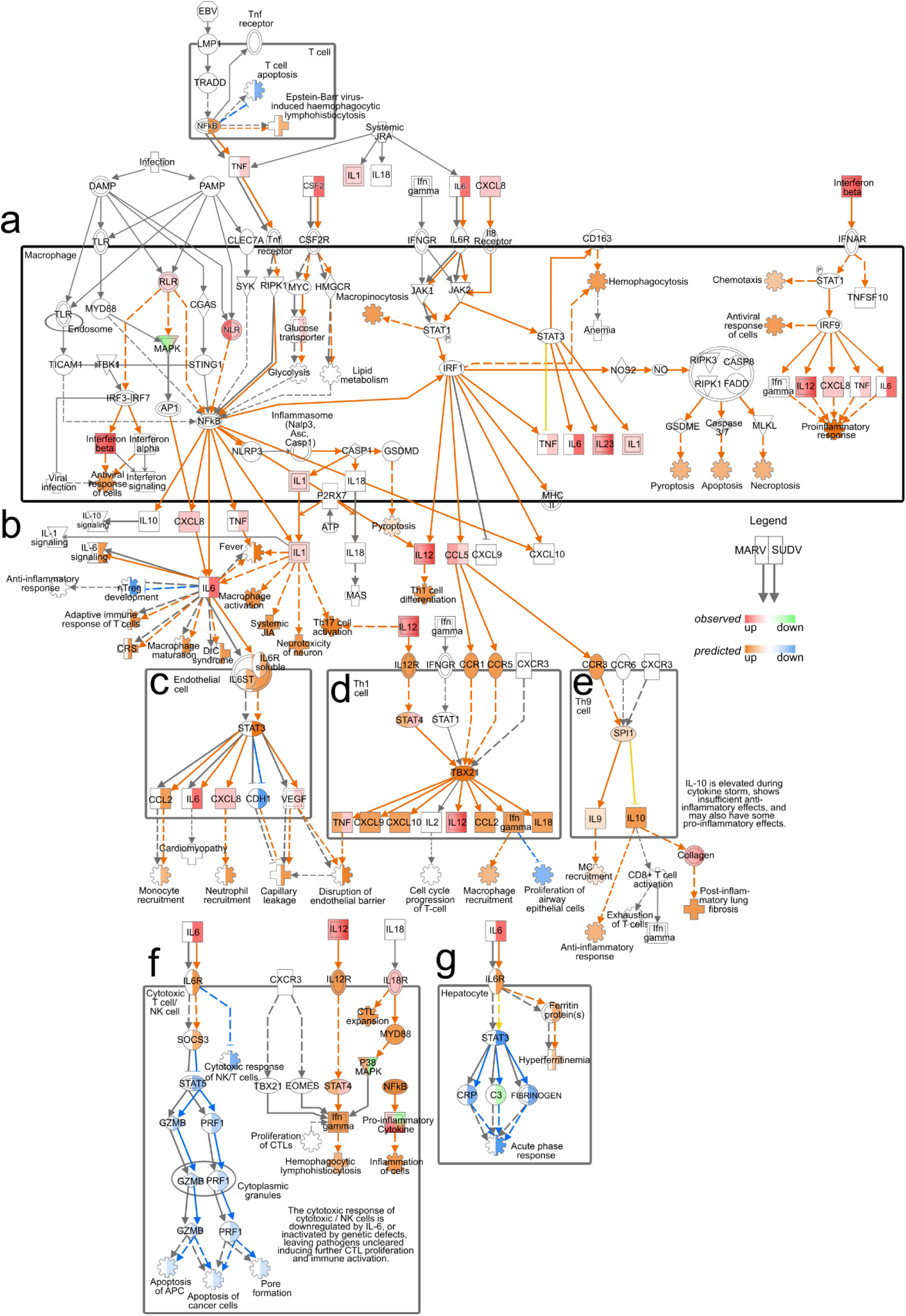
Pathogen Induced Cytokine Storm Signaling Pathway regulation by MARV and SUDV in ERBs. The map was generated using the IPA molecular activity predictor (MAP) analysis of the canonical Pathogen Induced Cytokine Storm Signaling Pathway (a) in macrophages, (b) exogenously, (c) in endothelial cells, (d) in Th1 cells, (e) in Th9 cells, (f) in cytotoxic T cells/NK cells and (g) in hepatocytes. Within macrophages and for exogenous gene expression (a-b), only genes actually observed in our data as differentially expressed are indicated as red (upregulated) or green (downregulated), while downstream responses (cartwheel and cross symbols) predicted as differentially regulated (orange for upregulated and blue for downregulated) based on the true DEGs are included. For the downstream signaling and responses in other cell types in this pathway (c-g) both real observed (red-green) and predicted (orange-blue) gene expression and downstream processes are included. For every symbol, the left half always corresponds to the relevant response or prediction for MARV, and the right half corresponds to the relevant response or prediction for SUDV. Orange arrows represent predicted upregulation, blue arrows predicted downregulation, yellow arrows ambiguous prediction and grey arrows no differential regulation.

In T cells, the predicted upregulation of *NFκB* in response to SUDV was forecasted to suppress T cell apoptosis, contrasting the absence of predicted *NFκB* expression for MARV-infected cells. Within the macrophage compartment, the observed upregulated expression of *TNF, IL6* and *CSF2* in response to SUDV, but not MARV, reflected in the differential prediction for *RIPK1* and *NFkB* signaling, glucose transport, lipid metabolism, glycolysis and JAK1/JAK2 signaling between MARV and SUDV (**Fig. 7a**). Despite the differential *TNF* and *IL6* responses, however, exogenous *IFNB* signaling and the upregulation of *IL12* and *CXCL8* in response to both viruses in macrophages resulted in similar predictions for upregulated chemotaxis, antiviral response of cells and proinflammatory response (**Fig. 7a**). Exogenously, the upregulated expression of IL6 signaling by SUDV was predicted to induce diverse downstream processes, including adaptive immune response of T cells, macrophage activation and maturation, fever and the suppression of nTreg development, none of which were predicted as induced in MARV-infected cells. In contrast, the upregulation of *IL12* in response to both viruses was predicted to induce Th1 cell differentiation **(Fig. 7b**).

The observed upregulated expression of *CXCL8, IL1, IL12* and *CCL5*, coupled with the divergent *TNF* and *IL6* responses were predicted to heavily influence the forecasted downstream signaling processes in diverse cell types. In endothelial cells, *IL6* signaling from SUDV-infected but not MARV-infected macrophages was predicted to induce *STAT3* expression, the upregulation of *CCL2* and the suppression of *CDH1*. Combined with the observed upregulation of *CXCL8* and *VEGF*, the predicted outcome for SUDV infection included increased monocyte and neutrophil recruitment, paralleled by capillary leakage and disruption of the endothelial barrier (**Fig. 7c**). In Th1 and Th9 cells, on the other hand, *CCL5* signaling through the chemokine receptors CCR1, CCR5 and CCR3 was predicted to induce comparable upregulated expression of *STAT4, TBX21, CXCL9, CXCL10, CCL2, IFNγ* and *IL18* in response to both viruses. Downstream macrophage and mast cell (MC) recruitment, the suppression of proliferation of airway epithelial cells, anti-inflammatory response and post-inflammatory lung fibrosis were also predicted as similarly regulated in MARV and SUDV-infected cells (**Fig. 7d-e**).

A strong and clear difference in cytokine signaling between MARV and SUDV was also predicted in cytotoxic T cells/NK cells and hepatocytes. In cytotoxic T cells/NK cells, we found that SUDV-induced *IL-6* was predicted to induce *SOCS3* expression and the downstream suppression of *STAT5*, granzyme B (*GZMB*) and *PRF1* expression, leading to suppressed apoptosis of APCs and cancer cells, as well as pore formation (**Fig. 7f**). In hepatocytes, on the other hand, the presence of *IL6* signaling in SUDV-infected cells was predicted to induce hyperferritinemia, the suppression of *STAT3*, C-reactive protein (*CRP*), fibrinogen expression and the downregulation of acute phase response (**Fig. 7g**). Due to the absence of *IL6* signaling in MARV-infected cells, these genes and pathways remained blank in the MAP analysis, indicating the absence of MARV-induced regulation along these signaling cascades in cytotoxic T cells/NK cells and hepatocytes (**Fig. 7g**).

### ERB bmMΦs display a distinct response to MARV compared with human MΦs

Finally, we compared the transcriptional responses of ERB MΦs observed herein against our recently published dataset of human monocyte-derived MΦ (moMΦ) responses to MARV to directly relate any similarities or differences in cell responses in the natural reservoir versus the spillover host^36^. For this, we first quantified and compared the intracellular viral replication in ERB and human moMΦs by calculating the percentage of the total gene counts that constituted viral genes in each species. We found very similar intracellular MARV RNA loads in each host, with viral gene counts constituting 1.09% of total gene counts in ERB bmMΦs and 0.96% in human moMΦs (**Fig. 8a, b**). We then compared viral loads in cell culture supernatants and found that despite similar MARV-NP gene copies intracellularly and in cell culture supernatants, human moMΦs released significantly more infectious viral particles than ERB bmMΦs (**Fig. 8b**).

**Figure 8.**
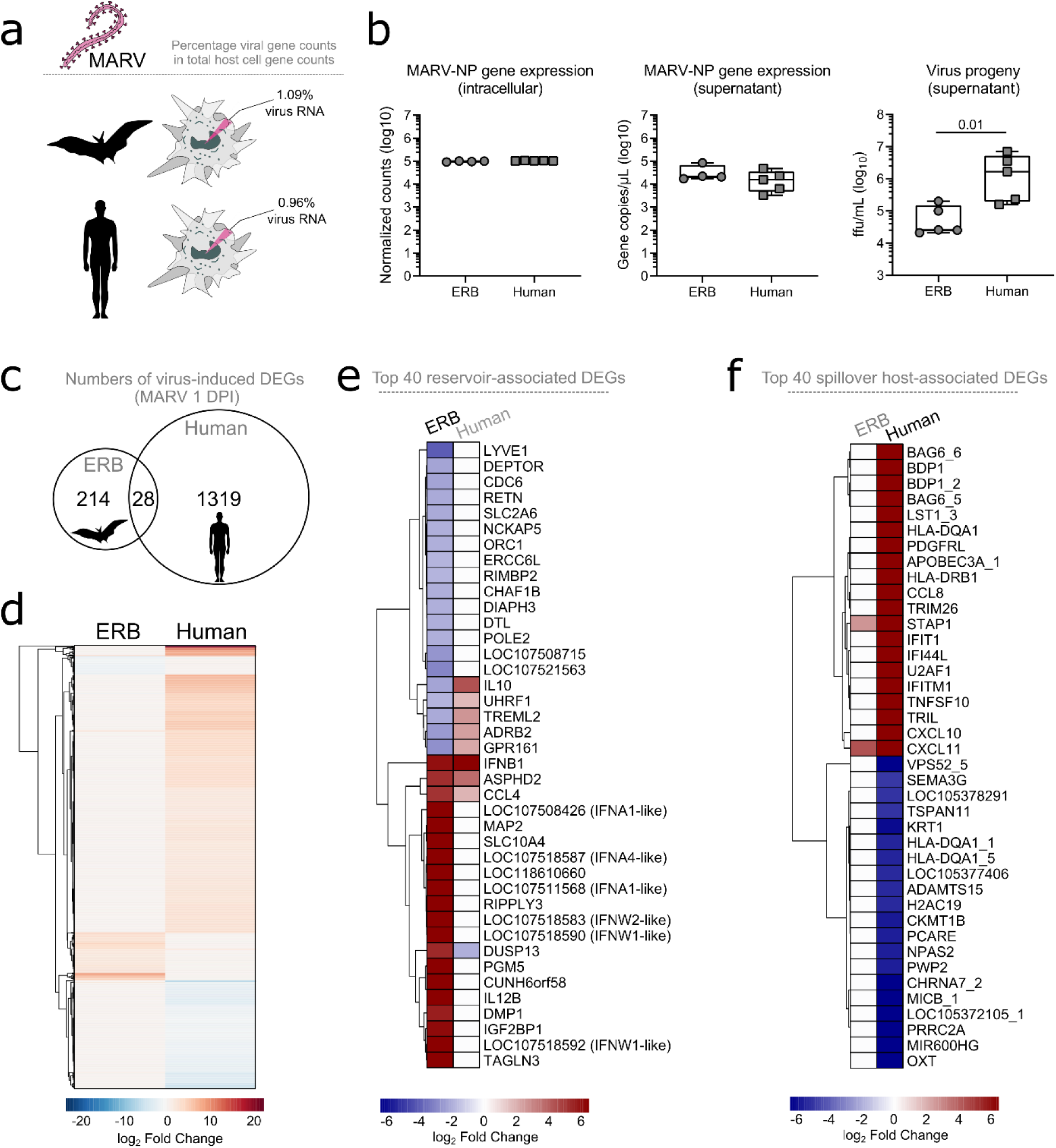
Comparative DEG profile of ERB-derived bmMΦs and human monocyte-derived MΦs (mo MΦs) at 1 DPI with MARV. (a) Percentages of viral RNA within total RNA counts in ERB and human MΦs. The percentage was calculated using the sum of normalized gene counts of all 7 MARV genes against the sum of all host gene counts within the bulk RNAseq dataset for each species. (b) Viral replication represented as intracellular MARV-NP gene counts (left), MARV-NP gene counts in cell culture supernatants (middle) and viral progeny in cell culture supernatants (right). (c) A Venn diagram illustrating the total numbers of unique and shared DEGs induced by MARV in ERB bmMΦs and human moMΦs at 1 DPI. (d) Global heatmap of DEG expression in MARV-infected ERB bmMΦs and human moMΦs at 1 DPI. (e) A heatmap of the top 20 upregulated and top 20 downregulated genes induced in MARV-infected ERB bmMΦs, plotted against the expression of their human orthologs. (f) A heatmap of the top 20 upregulated and top 20 downregulated genes induced in MARV-infected human moMΦs, plotted against the expression of their ERB orthologs. DEGs for each species were defined as genes with a padj < 0.05 and a log_2_ fold change ≥ ± 1.5. The ERB dataset includes pooled data from four individual bats, while the human dataset includes pooled data from three individual healthy donors. The heatmaps are shown as log_2_ fold change against respective mock-infected negative controls for each species.

The early transcriptional response to MARV included the differential expression of 1319 genes in human moMΦs, contrasted by 214 genes induced in ERBs and only 28 shared DEGs induced in both species, highlighting significant qualitative and quantitative differences in bat and human macrophage responses to MARV *in vitro* (**Fig. 8c, d**). To explore in greater detail the MΦ transcriptional profile in each species, we extracted from each dataset the top 20 up- and top 20 down-regulated genes of ERBs and humans, designating each gene set as the “Top 40 reservoir-associated DEGs” and “Top 40 spillover host-associated DEGs”, respectively (**Fig. 8e, f**). We observed very limited overlap in antiviral response gene expression between ERBs and humans, with only *ASPHD2, IFNB1* and *CCL4* being similarly upregulated in both hosts. The dual specificity phosphatase gene *DUSP13* was upregulated in ERBs, but downregulated in humans, while five other genes were downregulated in ERBs but upregulated in human moMΦs (*IL10, UHRF1, TREML2, ADRB2, GPR161*) (**Fig. 8e**). Interestingly, *MAP2, RIPPLY3* and *TAGLN3* shown earlier to be upregulated by MARV and not SUDV in ERBs were not induced in MARV-infected human moMΦs either, further indicating a specific role for these cytoskeleton and polymerase-associated genes restricted to the ERB-MARV context (**Fig. 4d**; **Fig. 8e**). In contrast, within the top 40 spillover host-associated DEGs, only two genes were mutually upregulated in both ERBs and humans – the chemokine *CXCL11* and the signal transduction adaptor protein gene *STAP1* (**Fig. 8f**). Unlike ERBs, human moMΦs also displayed simultaneous upregulation and downregulation of various HLA genes (*HLA-DQA1s* and *HLA-DRB1*), cytokines and chemokines (*CCL8, CXCL10, CXCL11, TNFSF10*), ISGs (*IFIT1, IFI44L, IFTM1*) and various antiviral-associated genes (*APOBEC3A, TRIM26*), classical signs of filovirus-induced deregulation of host macrophage functionality (**Fig. 8f**).

## Discussion

ERBs have been extensively established as natural reservoirs of MARV and while the reservoirs of pathogenic orthoebolaviruses like EBOV and SUDV are yet to be discovered, several bat species are considered credible candidates^18,19,39^. ERBs represent unlikely orthoebolavirus reservoirs, as they’re largely refractory to experimental infections with EBOV, BDBV, TAFV or RESTV^26,40^. However, SUDV is capable of limited tissue-restricted replication in ERB liver, spleen and axillary lymph nodes, offering an invaluable opportunity to explore what features of the ERB innate immune response are modulated through co-adaptation with MARV, in contrast with protective immunity to SUDV infection. More specifically, we sought to compare MΦ transcriptional responses to MARV and SUDV with the assumption that while MARV will induce unique transcriptional regulation driven by co-evolution with ERBs, SUDV should induce non-adapted responses resembling those of a foreign viral infection. This would allow us to elucidate some of the augmented features of the reservoir-virus response that allow for population-level maintenance of MARV, which likely accommodate sufficient MARV replication in an ERB to facilitate onward virus transmission to other bats, as opposed to being cleared within days with no significant shedding like that seen in SUDV-infected ERBs. We chose MΦs as they represent an important early host target of filoviruses in both humans and bats, their dysregulation being a central factor in primate filovirus disease progression and immunopathology.

Gaining a deeper understanding of the molecular and cellular mechanisms underlying the zoonotic reservoir competence of bats has been confounded by a distinct paucity of species-specific laboratory reagents and assays. Recent in-depth analyses of bat genomes have revealed new insights into numerous evolutionary adaptations of chiropteran immune systems that some viruses have likely leveraged as platforms for co-adaptation, contributing to the zoonotic reservoir status of some bats. Various bat species appear to have acquired expanded natural killer cell receptor genes^41^ or alternatively have undergone gene loss of *α* and *β defensin* genes^42^, killer cell lectin-like receptor K1 (*KLRK1*), PYRIN and HIN domain (*PYHIN*) genes^43,44^ or proinflammatory cytokine genes like *IL36A* and *IL36G*^9^. Others have experienced significant positive selection of viral sensors and inflammatory response regulators like *TLR8* and *TRIM38* or the deletion of function-altering Cys residues in IFN-associated genes like *ISG15*^9^. Unique patterns of IRF7-driven induction of ISGs in black flying foxes (*P. alecto*) were also recently reported^43^. Combined, these findings illustrate that different bats have evolved diverse and highly-specific molecular mechanisms that likely contribute to their ability to tolerate, maintain and transmit some viral pathogens by striking a fine-tuned balance between the induction of sufficient antiviral immune responses and the absence of aberrant tissue-damaging inflammatory processes.

Studies have previously reported the successful *in vitro* differentiation and characterization of bat bmMΦs from black flying foxes (*P. alecto*) and cave nectar bats (*Eonycteris spelaea*)^44,45^. However, few studies have focused on developing MΦ cultures from known bat reservoirs of zoonotic diseases. Taking advantage of limited bat resources, we successfully differentiated ERB-derived bmMΦs for an immunological interrogation of filovirus-specific *in vitro* responses. In line with the reservoir competence of ERBs, their bmMΦs were readily susceptible to MARV and maintained stable low-level virus replication over the course of 3 days, similar to ERB-derived bmDCs^28^. Moreover, the previously reported transcriptional profile of MARV-infected bmDCs reflected in our current observations of the upregulation of only a restricted set of cytokines, chemokines, Type I IFNs, ISGs and transcriptional factors in MARV-infected bmMΦs mostly restricted to 1 DPI^28^. For SUDV we detected notably higher intracellular viral loads than for MARV at the same MOI, coupled with declining viral production in cell culture supernatants, mirroring the *in vivo* differential control kinetics of MARV and SUDV reported following experimental infections of ERBs^26^.

Regardless of the immune or non-immune origin of ERB cells (bmMΦ vs. RoNi), MARV infected fewer cells and replicated less than SUDV over a 3-day *in vitro* infection, contrasting comparable replication rates of MARV and SUDV in Vero E6 cultures. Moreover, despite similar intracellular replication in bat and human MΦs, MARV-infected ERB bmMΦ cultures contained significantly fewer infectious viral particles than human moMΦs, indicating the presence of host-specific intrinsic differences in viral replication processes unique to the ERB-MARV relationship.

Viruses have evolved to rely on, exploit and remodel the host cell machinery for their entry, replication, trafficking, shuttling and virion egress^46^. As a result, they also inadvertently remodel the host cell actin cytoskeleton and can even prime RIG-I-like receptor activation in response to cytoskeleton disturbances^47^. The directed transport of viral proteins typically takes place alongside cytoskeletal tracks like microtubules or actin filaments^48–50^. Here, we found that the top DEGs induced by MARV in ERB bmMΦs included several genes encoding microtubule-associated proteins (MAPs) and two genes associated with RNA polymerase activity. Importantly, neither SUDV-infected bat bmMΦs, nor MARV-infected human moMΦs induced the expression of any of these genes, highlighting the presence of host cell cytoskeleton and polymerase activity alterations unique to MARV-ERB infection settings. Prior evidence of filovirus protein association with host cell microtubules has shown an association of EBOV matrix protein VP40 with host cell microtubules, resulting in the stabilization of cellular microtubules against drug-induced depolymerization and enhancement of tubulin polymerization - properties similar to those of MAPs^46^. In contrast, drug-induced depolymerization of microtubules in a human macrophage cell line, but not in other cell lines, increases MARV viral protein release *in vitro*, indicating a cell type-specific association between MARV viral progeny release and host cell microtubule organization^51^. Based on these and our own findings, we therefore hypothesize that the ability of MARV to maintain low-level productive infections in these bats could be aided by viral co-adaptations that manipulate the intracellular replication machinery of ERBs to preserve viral replication rates sufficient to maintain viral transmission, but low enough to avoid inducing overt immune responses and rapid viral clearance.

Despite the stable MARV replication rates at 1, 2 and 3 DPI, ERB bmMΦs underwent an almost complete transcriptional shut-down by 3 DPI, contrasting the stronger and more diverse transcriptional changes maintained in SUDV-infected cells at the same timepoints. This striking phenomenon was evident both when looking at immune-related DEG expression and in our IPA dataset, which highlighted the presence of MARV-induced suppression of canonical signalling pathways associated with cell cycle control, DNA synthesis, mitosis and cell senescence. Together, these findings suggest the presence of expansive transcriptional silencing of a wide range of cellular processes in MARV-infected ERB bmMΦs by 3 DPI. Moreover, even though MARV replicated at similar rates in bat and human MΦs, we observed stronger and more diverse transcriptional responses to MARV in human MΦs than ERB bmMΦs. Viral replication dynamics alone are therefore an unlikely driver of the magnitude and nature of the transcriptional response profile of filovirus-infected host cells described herein. These responses likely rely on a combination of complex factors involving species-specific virus-driven immune modulation and antagonism, and in the case of ERBs and MARV, the accumulation of numerous co-evolutionary adaptations in both the reservoir and virus.

The exceptionally high virulence of filoviruses in humans and NHPs is at least partially explained by the potent immune inhibitory properties of several virus-encoded proteins. Each filovirus employs its own strategies interfering with host antiviral immune responses. EBOV-VP24 inhibits IFN-induced JAK-STAT signalling^33^, while MARV-VP40 blocks STAT protein tyrosine phosphorylation^52^. EBOV-VP35 and MARV-VP35 both block IFN production by interfering with RIG-I signalling, albeit through different dsRNA interaction mechanisms and with different efficiencies^53–58^. Mutating specific MARV-VP35 residues associated with dsRNA binding results in improved Type I IFN responses and reduced viral replication, demonstrating a central role of VP35 as a virulence factor^53,54,59–61^. Herein, we found stark differences in MARV-VP35 and SUDV-VP35 correlation with several bat pattern recognition receptors (PRRs), Type I and III IFNs and ISGs. MARV-VP35 showed severely limited negative correlation with any of the tested DEGs, while SUDV-VP35 displayed moderate to strong negative correlation with most IFN-associated genes, indicating intact IFN antagonistic properties of SUDV-VP35. Contrasting MARV-VP35, MARV-VP40 showed a markedly stronger negative correlation with *RIG-I*, Type I and III IFNs, IFN receptors *IFNAR2* and *IL10RB*, *IRF7*, *NFKB1, ISG15* and *ISG20* than either SUDV-VP24 or SUDV-VP40. Considering the largely muted transcriptional changes observed in MARV-infected bmMΦs by 3 DPI, it is therefore conceivable that MARV-VP40 has likely co-evolved selective and unique IFN antagonistic properties that target ERB antiviral responses more efficiently than other filoviruses and contribute to the ability of these bats to maintain low-level MARV infections in the general absence of significant IFN-driven responses and rapid viral clearance. In contrast, our findings suggested that MARV-VP35 possibly displays reduced antagonistic abilities in these bats.

The type I (α, β, δ, ω and ε) and Type III IFN families (λ) represent key components of the early host antiviral immune response. The initial recognition of viral RNA by host PRRs induces the production of IFNs, which bind and signal through the heterodimeric receptor complex IFNAR1/2 (Type I IFN) or IFNLR1/IL10RB (Type III IFN) and trigger the downstream expression of diverse ISGs and in the case of IFNλs, also the induction of B- and T-cell driven adaptive immune responses^62–67^. Studies have now identified various noteworthy differences in the IFN repertoires of humans and bats, such as a contracted Type I IFN locus in black flying foxes (*P. alecto*) or an expansion of Type I IFN loci in large flying foxes (*P. vampyrus*) and little brown bats (*Myotis lucifugus*)^68,69^. In ERBs, the IFN-ω gene family has undergone considerable expansion and contains 22 *IFNW* genes, contrasting only 5 and 6 *IFNW* genes in *P. alecto* and *Rhinolophus ferrumequinum* bats, respectively, and a single *IFNW* gene in humans^41,70^. Even though *IFNW* genes don’t show constitutive expression in ERBs, SeV infection induces upregulated transcript levels in immortalized RoNi cells^41^. Moreover, recombinant IFN-ω4 can block experimental infection with a recombinant VSV in the same cell line, demonstrating measurable antiviral activity of ERB-derived IFN-ω^41^. Herein, we found that infection with MARV and SUDV induced an overall similar pattern of upregulated Type I IFN expression. However, MARV failed to induce type III IFN transcription in bmMΦs, while SUDV triggered the strong upregulated expression of two Type III IFN genes (*IFNL1-like* and *IFNL3)* at 2 and 3 DPI. Interestingly, ERB-derived lung organoids undergo significant upregulation of both the *IFN1-like* and *IFNL3* genes at 3 days post-infection with MARV, as well as in response to SeV, H1N1 influenza virus and VSV infections. Moreover, this Type III IFN response in bat organoids appears to drive robust, protective and self-amplified antiviral responses^71^. Considering the strong negative correlation between MARV-VP40, *IL10RB, IFNL1-like, IFNL3*, various transcriptional factors and ISGs, we therefore hypothesize that MARV has evolved unique VP40 antagonistic properties that specifically target ERB Type III IFN responses, subduing IFNλ production, downstream signalling and adaptive immune response activation. Additionally, given the key role of macrophages as both early targets of filoviruses and coordinators of innate and adaptive host responses, the absence of Type III IFN responses in ERB-derived MΦs could be a cell-specific response aimed at subduing downstream activation of further adaptive immune responses to MARV. In contrast, the weaker SUDV-VP24 and SUDV-VP35 negative correlation with these genes indicates that unlike MARV, SUDV infection induces sufficient Type III IFN responses in ERBs to potentially contribute to increased T and B cell proliferation and the ability of these bats to rapidly control SUDV infections *in vivo*.

To test whether the overall pathogen response observed in ERB bmMΦs potentially translates in differential adaptive immune responses to filoviruses, we performed a comprehensive IPA analysis of the canonical *Pathogen Induced Cytokine Storm Signaling Pathway*, regulated by both MARV and SUDV in these cells. We found that the absence of *TNF* and *IL6* signaling in MARV-infected cells and the significant upregulation of both cytokines in SUDV-infected cells lead to significant disparities in downstream immune cell activation and regulation. In response to SUDV, *TNF* and *IL6* signalling were predicted to suppress T cell and APC apoptosis, Treg development, pore formation and acute phase response in hepatocytes. The *TNF* and *IL6* signalling cascades were predicted to induce adaptive immune responses of T cells, the recruitment of various innate immune cells, including monocytes, macrophages, neutrophils and mast cells, as well as both antiviral responses and anti-inflammatory responses. Combined with the induction of Type III IFNs by SUDV and not MARV, our findings therefore highlight starkly different responses of ERBs to each virus and point to the induction of an unperturbed and well-balanced proinflammatory response to SUDV, paralleled by downstream recruitment and activation of endothelial cells, hepatocytes and T cells that likely contribute to the previously reported clearance of SUDV infection *in vivo*^26^. In contrast, the absence of *TNF, IL6* and Type III IFN responses to MARV in these bmMΦs are likely contributing factors to the muted transcriptional response to MARV in these cells. Unlike ERBs, elevated TNF, IL-6 and IL-10 cytokine responses are classical hallmarks of severe filovirus disease in both humans and NHPs following natural exposure or experimental infections^72–76^. Exposing human or NHP peripheral blood mononuclear cells (PBMCs) to filoviral peptides or inactivated viral particles also results in cell apoptosis, inhibition of CD4 and CD8 T cell cycle and maturation, and increased IL-10 production, resulting in an overall dysfunctional T cell response and the development of severe tissue pathology^77^. This pronounced immune dysfunction appears to be long-lasting, with EBOV survivors displaying elevated blood markers of inflammation, including high levels of IL-1β, TNF and CCL5, increased anti-inflammatory IL-10, sustained T cell activation and DC depletion 19-25 months post-infection, none of which are evident in filovirus-infected ERB bmMΦs^78^.

Additional work in our group recently offered a comprehensive comparative peripheral blood response analysis of ERB and NHP/human responses to MARV, EBOV and SUDV and highlighted remarkable consistency in transcriptional responses to all three viruses across primate studies^79^. Despite marked differences in experimental set-ups between these studies, a core set of canonical genes typically associated with mammalian antiviral responses and pathogenesis were evident in humans, NHPs and bats^79^. Those included key PRRs (*IFIH1/MDA5* and *DDX58/RIG-I*) and antiviral genes (eg. *ISG15, ISG20, IRF7, MX1, OAS1, OAS3, IFITs, STAT1, STAT2, FOS*). We also described clear divergent peripheral immune responses to filoviruses between bats and primates, the latter significantly upregulating various recognition receptors (eg. *TLR3, TLR4, DHX58*), antiviral genes (eg. *IFI35, OAS2, MX2*), proinflammatory cytokine and chemokine genes (eg. *CXCL10, IL6, CCL2/3/8, CXCL11, IL1B*), in line with the highly activated proinflammatory transcriptional profile of MARV-infected human macrophages^36,79^. As natural reservoirs, ERBs likely hold a number of critical evolutionary advantages in modulating MARV infection, replication and transmission over primate spillover hosts, whilst remaining asymptomatic and avoiding overt proinflammatory processes.

Herein, we found that beyond bulk blood cell or tissue-level responses, ERBs employ carefully fine-tuned pathogen-specific responses to different filoviruses at specific innate immune cell levels. Bat bmMΦ responses were characterized by a muted antiviral response to MARV, contrasted by a stronger, sustained and proinflammatory-skewed response to SUDV reminiscent of the strong filovirus-induced responses in humans and NHPs. Despite the presence of limited shared gene signatures between MARV and SUDV responses in ERBs, and even fewer similarities between bat and human responses to MARV, we identify several molecular mechanisms differentially regulated in these bats. We show evidence of virus-specific host cell cytoskeletal changes, unique patterns of viral protein antagonism, type III IFN responses, as well as differential TNF and IL6 responses. The absence of these mechanisms in response to MARV are a possible result of the highly specific coevolutionary relationship between MARV and its natural wildlife reservoir, allowing these bats to maintain and transmit MARV at low levels without developing signs of viral hemorrhagic fever disease themselves. In contrast, their induction in response to other filoviruses is a likely contributing factor to the ability of ERBs to clear orthoebolaviruses like SUDV. Even though these bats control MARV infections in the wild, they do allow for sufficient viral replication and persistence to maintain and transmit MARV at the population level. Thus, based on our findings, MARV has likely co-adapted to ERBs in a way that tempers innate immune responses enough to allow low-level viral replication sufficient for transmission in the absence of aberrant innate immune cell activation, but also without interfering with the generation of protective T and B cell responses.

## Methods

### *In vitro* differentiation of ERB-derived bmMΦs

Bone marrow cells were obtained from captive ERBs euthanized for unrelated studies at the CDC following cell isolation protocols as described by Prescott et al. (2019)^28^. To differentiate bmMΦs, one vial of cryopreserved bone marrow cells per bat was thawed and resuspended in 9 mL of R10 medium containing 10% FCS, 1% L-glutamine, 1% penicillin and streptomycin, 1% HEPES and benzonase (10 µL/100mL medium) in RPMI-1640 medium (Sigma). The cell suspension was centrifuged at 350x g for 10 minutes and the R10 wash medium was carefully removed. The cell pellet was slowly resuspended in fresh R10 medium containing 20 ng/mL recombinant ERB macrophage colony-stimulating factor (rM-CSF, Kingfisher Biotech). The cells were then plated out at an approximate density of 5×10^5^ cells/well in a final volume of 250 µL and incubated at 37°C in 5% CO_2_. After one day of incubation, the cells were supplemented with 250 µL/well of fresh pre-warmed R10+M-CSF medium. The medium was added slowly and drop-wise to the center of each well. The plate was then returned to the incubator. On days 3 and 6, half of the medium in each well was carefully removed and was replaced with 250 µL of fresh pre-warmed R10+M-CSF medium as described above.

On day 8 of differentiation, the cell culture medium was carefully removed, and adherent cells were gently washed by slowly adding 500 µL/well of pre-warmed PBS and carefully removing it again. At this stage, cell density and morphology were controlled visually under a microscope. One well of cells per bat on each plate was always designated for cell dissociation and counting prior to restimulation or infection.

### Cell stimulation and virus infections

Prior to infection or restimulation, the culture medium in the wells containing bmMΦs designated for cell counting was carefully removed. The cells were washed with 500 µL/well of pre-warmed PBS as described above and Cell Dissociation Buffer (Life Technologies Corporation) was added to each well. In brief, for cell dissociation, 500 µL of Cell Dissociation buffer was added to each well, and cells were incubated at room temperature for 15 min, occasionally tapping and swirling the plate to facilitate cell detachment from the plastic. After 15 min, the cells were then gently dissociated by repeated pipetting and scraping of each well with the pipette tip. The buffer containing the detached cells was transferred in fresh 2 mL centrifuge tubes. The wells were washed with 500 µL of PBS, which was then added to the cell suspension. The cells were centrifuged at 350x g for 5 minutes. The buffer was then carefully removed and the cells were resuspended in 500 µL of R10 medium. Dissociated cells were stained with trypan blue and were manually counted under a microscope using Neubauer chambers. The obtained cell counts were then used to calculate the appropriate volume of virus needed to achieve the desired multiplicity of infection (MOI).

For stimulation with bacteria lipopolysaccharide (LPS), bmMΦs were incubated in 250µL of R10+M-CSF medium containing 2 µg/mL LPS (InvivoGen). For Sendai virus (SeV) infection, cells were incubated in 250µL R10+M-CSF medium containing 30 hemagglutination (HA) units of the Cantell strain, a non-pathogenic paramyxovirus used as a positive stimulation control for host IFN signaling. For filovirus infections, bmMΦs were covered in 100 µL of R10+M-CSF medium to prevent desiccation and were infected with either MARV (isolate Uganda 200704852 Uganda Bat, MARV371), MARV expressing green fluorescent ZsG protein (MARV-ZsG), SUDV-Gulu or SUDV-Gulu expressing ZsG (SUDV-ZsG) at an MOI of 2 (as titrated on Vero E6 cells). Cells were incubated with each virus inoculum for 1 h at 37°C in 5% CO_2_ with gentle mixing every 15 min by slowly swirling the plate to ensure even inoculum distribution in each well. The virus inoculum was then carefully removed, the cells were washed in pre-warmed PBS and fresh 250µL R10+M-CSF medium was slowly added to each filovirus-infected well. Mock-infected bmMΦs were cultured in R10+M-CSF medium only. At indicated timepoints, cell culture supernatants were collected for virus isolation, viral and cell RNA extraction, while bmMΦs were dissociated from the plates as described above for staining and flow cytometry.

Immortalized RoNi cells and VeroE6 cells were plated out in 24-well tissue culture-treated plates at an approximate density of 2×10^5^ cells/well in 1mL/well of DMEM medium containing 10% FCS, 1% L-glutamine and 1% Penicillin/Streptomycin. In the BSL-4 lab, the medium was carefully removed using a multi-channel pipette and the cells were washed once using fresh pre-warmed DMEM medium. Each cell type was then infected with either MARV-ZsG or SUDV-ZsG at an MOI of 2. Mock-infected cells were included as controls. The protocol for cell infection, incubation, cell dissociation and staining for flow cytometry was as described above for ERB bmMΦs.

Work with wild-type and recombinant ZsG filoviruses was conducted at the Robert Koch Institute under Biosafety Level 4 (BSL-4) laboratory conditions. Research staff involved in this study adhered closely to all approved BSL-4 safety protocols and standard operating procedures (SOPs) for sample inactivation and removal from the BSL-4 facility.

### Flow cytometry

For flow cytometry analysis of bmMΦ surface marker expression, mock-infected or filovirus-infected cells were harvested as described above. Cells were transferred in 2mL tubes and were stained in 30µL per sample of antibody mix in FACS buffer (protein-free PBS containing 0.2% BSA and 2mM EDTA) with Live/Dead Fixable Yellow Dead Cell Stain Kit (Invitrogen) and antibodies raised against the following markers: anti-mouse CD11b-PE (clone M1/70 diluted 1:100, BD), anti-human HLA-DR-A785 (clone L243 diluted 1:50, BioLegend), anti-human CD40-PE-Cy7 (clone 5C3 diluted 1:20, BioLegend), anti-human CD163-AF674 (clone QA19A16 diluted 1:100, BioLegend) and anti-human CD206-PB (clone 15-2 diluted 1:100, BioLegend). A custom-made anti-bat CD14 antibody conjugated in-house with either PerCP or AF647 LightningLink kits (Abcam) as per the manufacturer’s instructions was also included in the staining panel (CDC, diluted 1:100). Cells were stained for 15 min at room temperature and were then washed once in 200µL/sample of FACS buffer. Stained and washed cells were fixed overnight in 200µL/sample of 10% formalin. Following overnight fixation, cells were transferred in fresh 200µL formalin and were removed from the BSL-4 laboratory in accordance with approved SOPs. Samples were run on a Cytoflex S cytometer (Beckman Coulter GmbH) and the final results were analyzed using FlowJo software version 10.8.1 (TreeStar).

### Confocal microscopy

For confocal fluorescence microscopy imaging, cells were fixed with 10% formalin (HistoFix, Roth) and were then stained with Acti-Stain 670 (Cytoskeleton) and DAPI (RotiMount, Roth) according to the manufacturer’s instructions. Imaging was performed using the Stellaris 8 confocal microscope (Leica) at the Unit for Advanced Light and Electron Microscopy, Center for Biological Threats and Special Pathogens at the Robert Koch Institute. Image processing was performed using ImageJ software.

### Real-time quantitative PCR (qPCR)

Viral RNA levels in cell culture supernatants were measured using real-time quantitative PCR. In brief, 140 µL of cell culture supernatant were collected per sample from mock-, MARV- and SUDV-infected bmMΦs at indicated timepoints of infection and were added to 560 µL of AVL buffer (Qiagen). For sample inactivation, 560 µL of 100% ethanol was added to the sample-AVL mix for removal from the BSL-4 facility following approved SOPs by trained personnel. RNA from these samples was extracted using the QIAamp Viral RNA Kit (Qiagen) following the manufacturer’s instructions. MARV and SUDV transcripts were quantified using a qPCR assay targeting the NP gene of each virus using an AgPath-ID One-Step RT-PCR Kit (Thermo Fischer Scientific). 25 µL reactions were formulated by adding 5 µL of sample into a master mix containing 10 µM of forward and reverse primers, 10 µM of TaqMan probe, 1x buffer, and 1x RT-PCR enzyme mix. The thermal profile used a 15 min incubation at 45°C, a 10 min incubation at 95°C, and 45 cycles of 15 s at 95°C, followed by 60 s at 60°C. Sample CT values for each virus were compared to a standard curve using MARV or SUDV transcripts of known concentrations ranging from 10^1^ to 10^6^ copies. Viral gene copies per µL cell culture supernatant were then calculated based on the standard curves. The primer and probe sequences used in this study are provided in Table S1.

### Gene expression analysis

For bulk RNA sequencing (RNAseq) of mock-infected, LPS-treated and virus-infected bmMΦs, adherent cells were lysed in 350 μL/well of RLT buffer. The cell-RLT mixture was then transferred in clean 2 mL sample tubes and were inactivated by adding 600 μL of 70% ethanol to the sample-RLT mix. Following inactivation, samples were removed from the BSL-4 facility by trained scientific staff following approved SOPs. Total RNA was extracted using the QIAGEN RNeasy Mini Kit following the manufacturer’s instructions. Extracted RNA samples were submitted to Novogene for library preparation, quality control and sequencing. In brief, messenger RNA (mRNA) was purified from total RNA using poly-T oligo-attached magnetic beads. Following fragmentation, the first strands of complementary DNAs (cDNA) were synthesized using random hexamer primers, followed by the second cDNA strand synthesis, end repair, A-tailing, adapter ligation, size selection, amplification, and purification. After final quality control, cDNA libraries were sequenced on multiple lanes using an Illumina NovaSeq platform.

The final RNAseq reads underwent quality control using FastQC^80^. Index adaptors were trimmed using Trim Galore and low-quality base-calls or reads below 20 base pairs were removed using a read quality cutoff Phred score of 33^81^. Trimmed quality-controlled reads were merged into a single file for each sample and were aligned against the *R. aegyptiacus* mRouAeg1.p reference genome (GenBank accession number GCA_014176215.1). For viral gene counts, trimmed and filtered reads were aligned against either the MARV ViralProj15199 (GenBank accession number GCF_000857325.2) or the SUDV ViralProj15012 (GenBank accession number GCF_000855585.1) reference genome. Gene level counts were quantified using Kallisto^82^, followed by filtering and log_2_ normalization of gene counts using the Tidyverse, BaseR and EdgeR packages in RStudio^83,84^. Differential gene expression analysis was performed using the Bioconductor package DESEQ2 to identify genes differentially expressed between mock-infected, LPS-treated and virus-infected bmMΦs^85^. Differentially expressed genes (DEGs) were defined as having a padj value < 0.05 and a log_2_ fold change expression of >1.5 for upregulated or <-1.5 for downregulated genes.

### Correlation analysis of gene expression

To assess the possibility of differential host immune response antagonism by viral proteins with known immunosuppressive properties, the log_2_ fold change values of MARV-VP35, MARV-VP40, SUDV-VP24 and SUDV-VP35 were extracted in a separate table. The complete list of DEGs was then manually inspected to identify and extract into the same table the log2 fold change values of all Type I and Type III IFN response-associated genes differentially expressed in virus-infected ERB at days 1, 2 and 3 post-infection. The complete table containing all viral protein and IFN gene expressions across the three timepoints was loaded in RStudio. Spearman’s rank correlation coefficient analysis was then performed for every gene against every other gene separately for MARV-infected and SUDV-infected samples using the corrr and corrplot RStudio packages. Significance levels were set at p<0.05 and insignificant correlations were blanked out the pyramid tables using corrplot.

### Ingenuity Pathway Analysis

Significantly enriched pathways and upstream regulators were determined using Ingenuity Pathway Analysis (IPA, QIAGEN Digital Insights, Redwood City, CA, USA) for each timepoint for SUDV and MARV. Datasets uploaded included DEGs and expression values with adjusted p values at each time point and analyzed using “Core Analysis” with default settings. Subsequent “Comparison Analysis” with default settings was performed to find commonalties in pathway enrichment and upstream regulators across timepoints for both viruses. Canonical pathways are ranked using a z-score algorithm that is calculated based upon dataset correlation from uploaded DEG and p-values with an activated state in that canonical pathway. P values result from a Fischer’s exact test that calculates the probability that the association between the genes in the uploaded dataset and the genes in the canonical pathway are due to chance alone. The pathway image was modified from the “Pathogen Induced Cytokine Storm Signaling Pathway” figure by removing downstream connections not relevant to the study. The heatmaps of canonical pathway and upstream regulator expression were created using GraphPad Prism 9 software (CA, USA).

### Statistical analysis

Log_2_ fold change values of significant DEGs following bulk RNA sequencing were based on Wald tests for differential expression and were defined as having a padj value < 0.05 and a log_2_ fold change expression of >1.5 for upregulated or <-1.5 for downregulated genes (Love et al., 2014). Gene expression correlation analyses were performed using the non-parametric Spearman’s rank correlation coefficient test, with a significance cut-off value set at p<0.05.

### Data availability

All data in this study are available from the corresponding author upon reasonable request. The code scripts used for the gene expression analysis used throughout the study are available on GitHub (https://github.com/ivetyorda/ERB_bmMac_MARVvsSUDV).

## Acknowledgements

This study received financial support through intramural funding from the Robert Koch Institute awarded to J.B.P. and through pilot project grant 01KI2210 of the German Ministry of Education and Research (BMBF) awarded to I.A.Y. L.T.A. is an Additional Ventures Catalyst to Independence Fellow and Bladder Cancer Advocacy Network Career Development Awardee. The funding bodies played no role in the study design, data collection, analysis, interpretation or in the preparation of the manuscript. The authors thank the Viral Special Pathogens Branch of the Centers for Disease Control and Prevention (CDC, USA) for providing the bone marrow samples used in this study, as well as Markus Kainulainen and César Albariño for providing the recombinant MARV-ZsG and SUDV-ZsG viral isolates. The findings and conclusions in this article are those of the authors and do not necessarily represent the official positions of the Centers for Disease Control and Prevention.

## Author contributions

I.A.Y. and J.B.P. conceptualized the study. I.A.Y. and A.L. performed experiments, acquired and analyzed the data. C.E.A. performed the Ingenuity Pathway Analysis (IPA). C.E.A. and L.T.A. provided bioinformatics support. N.C. performed the confocal microscopy analysis and J.C.G. contributed with the generation of ERB-derived bone marrow cells. J.S.T. and J.B.P. supervised the study. I.A.Y., C.E.A., N.C. and J.B.P. wrote the manuscript, and all authors revised its final version.

