## Supplementary Figures for "Egyptian rousette bat macrophages elicit divergent interferon responses and cytokine storm signalling against Marburg and Sudan viruses"

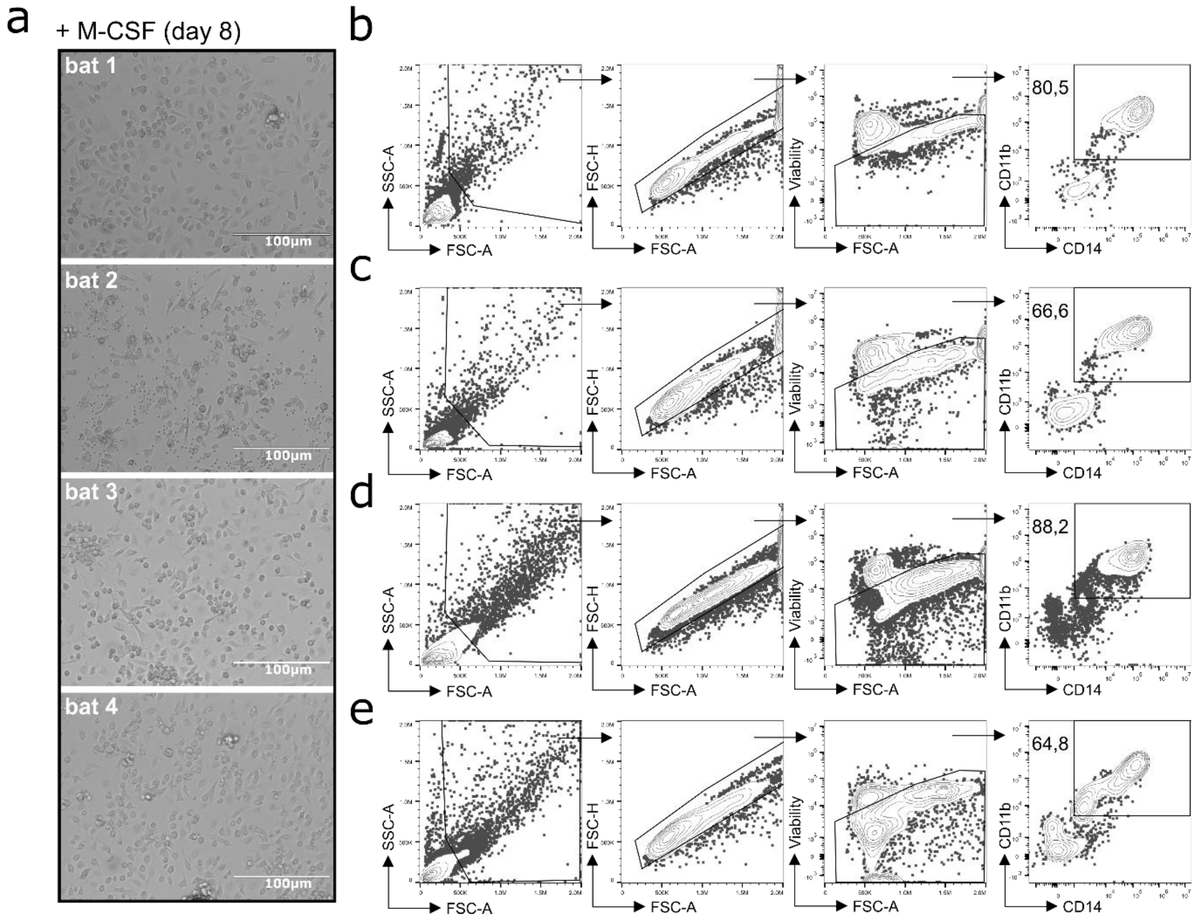


**Supplementary Figure 1.** Differentiation of Egyptian rousette bat (ERB) bone marrow-derived macrophages (bmMΦs). (a) Example light microscopy images illustrating the morphology of bone marrow cell cultures after 8 days of *in vitro* differentiation with recombinant ERB-specific M-CSF using cells derived from four individual bats. (b-e) Corresponding gating strategies and example contour plots showing the identification of $CD11b^{+}CD14^{+}$ bmMΦs via flow cytometry in the four individual bats.

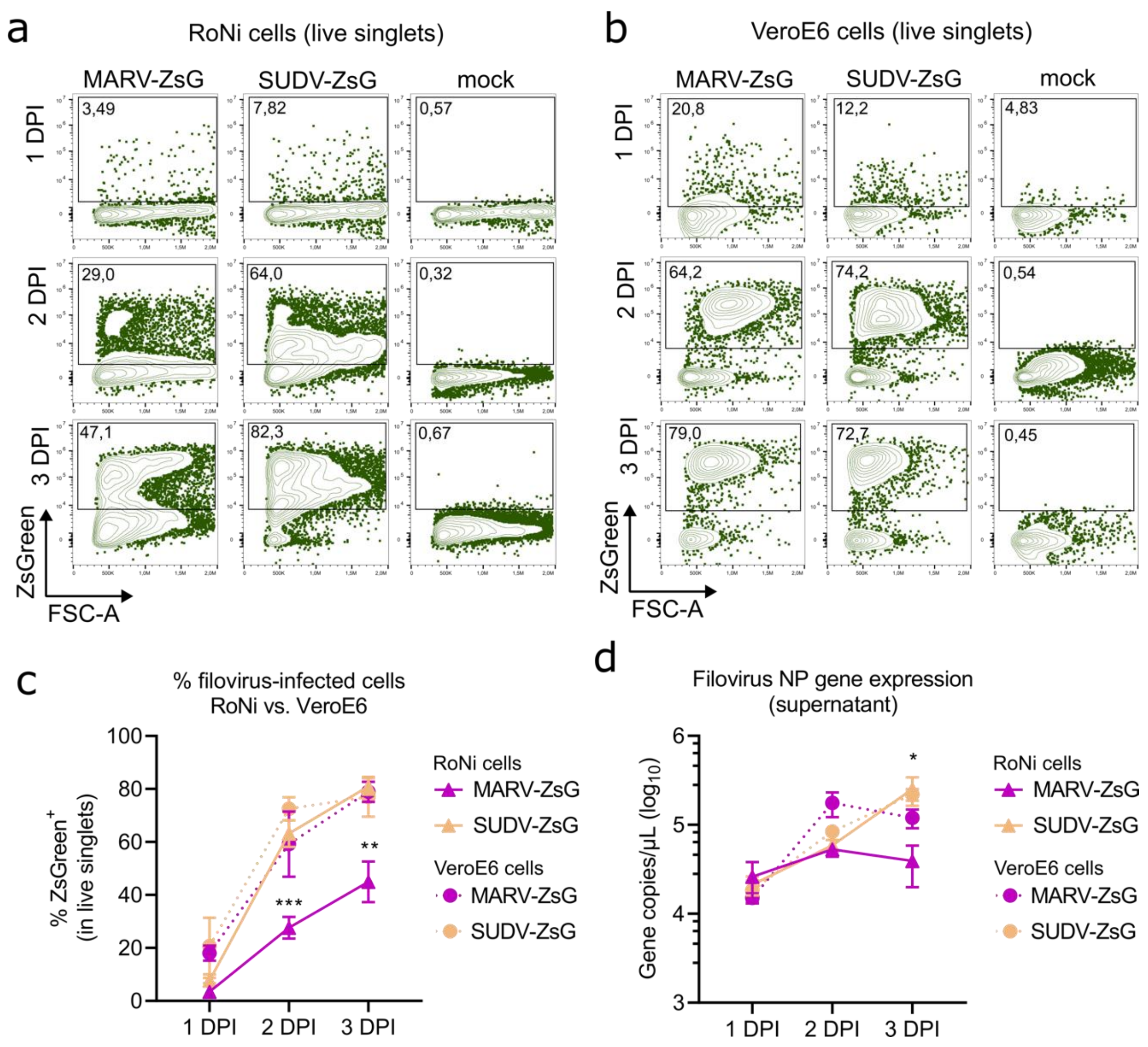


**Supplementary Figure 2.** Filovirus replication in RoNi and VeroE6 cell lines. Example contour plots of ZsGreen signal in live singlets in (a) RoNi cells and (b) VeroE6 cells infected with either MARV-ZsG or SUDV-ZsG, measured on a flow cytometer at 1, 2 and 3 DPI. (c) The percentage ZsGreen$^{+}$ cells in live singlets and (d) Filovirus NP gene copies in cell culture supernatants of MARV-ZsG and SUDV-ZsG infected RoNi and VeroE6 cells at 1, 2 and 3 DPI. Statistical analysis in (c) and (d) was performed using an unpaired student's t-test. Statistical significance stars mark significant differences between MARV-ZsG and SUDV-ZsG in RoNi cells. *p<0.05, **p<0.01, ***p<0.001.

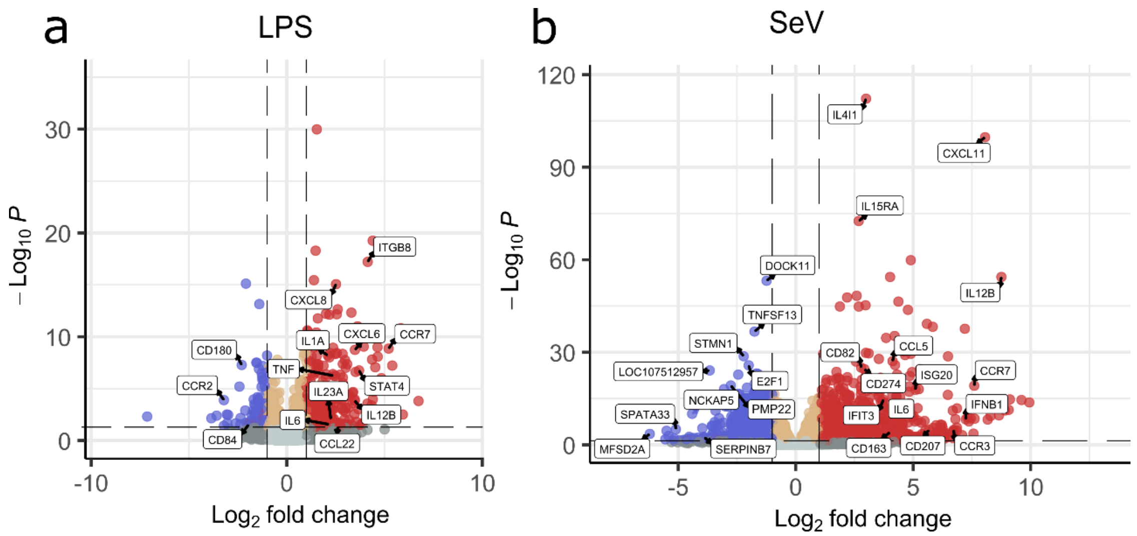


**Supplementary Figure 3.** ERB-derived bmMΦ responses to general immune stimulation. Volcano plots of significant DEGs across four individual bats in (a) LPS-stimulated and (b) SeV-infected bmMΦs, compared with mock-infected controls. DEGs were defined as genes with a padj $< 0.05$ and a $\log_2$ fold change $\geq \pm 1.5$. Genes with $\log_2$ fold change $\geq 1.5$ are marked in red (upregulated DEGs), genes with $\log_2$ fold change $\geq -1.5$ are marked in blue (downregulated DEGs). Genes with a $\log_2$ fold change $< \pm 1.5$ are marked in yellow, while genes with a padj $> 0.05$ are marked in grey (non-DEGs).

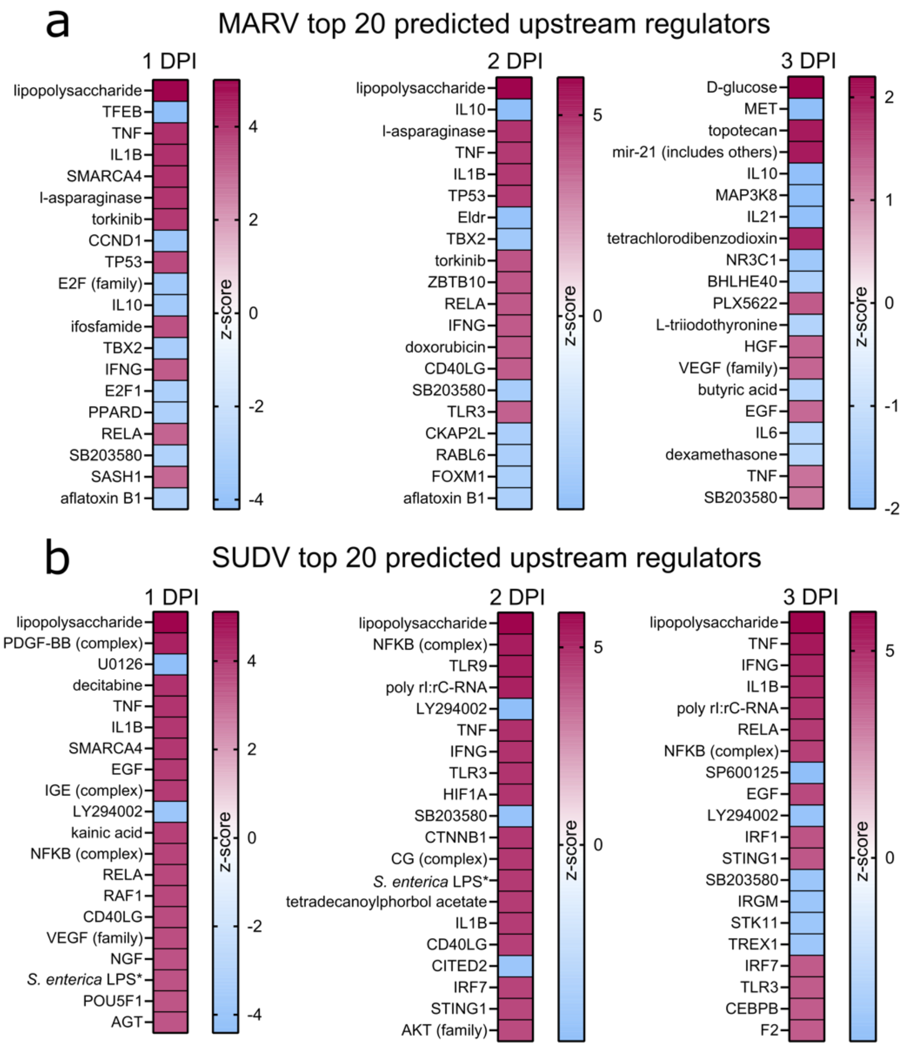


**Supplementary Figure 4.** Differential regulation of signaling pathways in filovirus-infected ERB bmMΦs. Ingenuity Pathway Analysis (IPA) of the statistically significant ($p<0.05$) (a) Top 20 predicted upstream regulators upregulated (magenta) or downregulated (blue) in response to MARV at 1, 2 and 3 DPI. (b) Top 20 predicted upstream regulators upregulated (magenta) or downregulated (blue) in response to SUDV at 1, 2 and 3 DPI.

**Supplementary Table 1.** Real-time quantitative PCR primer and probe sequences for quantification of viral RNA copies in cell culture supernatants, targeting MARV-NP and SUDV-NP.

| **Virus** | **Forward primer** | **Reverse primer** | **TaqMan probe** |
|---|---|---|---|
| MARV | GTCCTCAGCCAGAAACGAGA | ACCGTTACTTCCACAGGTGT | 6Fam-TCACAGAATCGGGTGTCACAGTCGT-BBQ |
| SUDV | GGTGGTGTTGTTGACCCGTA | CATCGTCGTCGTCCAAATTGA | 6Fam-TGAAGGCACCACAGGAGATCTTGATCT-BBQ |
